# Identification of an antiretroviral small molecule that appears to be a host-targeting inhibitor of HIV-1 assembly

**DOI:** 10.1101/2020.03.18.998088

**Authors:** Jonathan C. Reed, Dennis Solas, Anatoliy Kitaygorodskyy, Beverly Freeman, Dylan T. B. Ressler, Daryl J. Phuong, J. Victor Swain, Kent Matlack, Clarence R. Hurt, Vishwanath R. Lingappa, Jaisri R. Lingappa

## Abstract

Given the projected increase in multidrug resistant HIV-1, there is an urgent need for development of antiretrovirals that act on virus life-cycle stages that are not targeted by antiretrovirals currently in use. Host-targeting drugs are of particular interest because they can offer a high barrier to resistance. Here we report identification of two related small molecules that inhibit HIV-1 late events, a stage of the HIV-1 life cycle for which potent and specific inhibitors are lacking. This chemotype was discovered using cell-free protein synthesis and assembly systems that recapitulate intracellular host-catalyzed viral capsid assembly pathways. These compounds inhibit replication of HIV-1 in human T cell lines and PBMCs and are effective against a primary isolate. They reduce virus production, likely by inhibiting a post-translational step in HIV-1 Gag assembly. Notably, the compound colocalizes with HIV-1 Gag *in situ*; however, unexpectedly, selection experiments failed to identify compound-specific resistance mutations in *gag* or *pol*, even though known resistance mutations developed in a parallel nelfinavir selection. Thus, we hypothesized that instead of binding to Gag directly, these compounds might localize to assembly intermediates, the intracellular multiprotein complexes containing Gag and host factors that are formed during immature HIV-1 capsid assembly. Indeed, imaging of infected cells showed colocalization of the compound with two host enzymes found in assembly intermediates, ABCE1 and DDX6. While the exact target and mechanism of action of this chemotype remain to be determined, these findings suggest that these compounds represent first-in-class, host-targeting inhibitors of intracellular events in HIV-1 assembly.

**IMPORTANCE:** The success of antiretroviral treatment for HIV-1 is at risk of being undermined by the growing problem of drug resistance. Thus, there is a need to identify antiretrovirals that act on viral life cycle stages not targeted by drugs in use, such as the events of HIV-1 Gag assembly. To address this gap, we developed a compound screen that recapitulates the intracellular events of HIV-1 assembly, including viral-host interactions that promote assembly. This effort led to identification of a new chemotype that inhibits HIV-1 replication at nanomolar concentrations by inhibiting virus production. This compound colocalized with Gag and two host enzymes that facilitate capsid assembly but resistance selection did not result in compound-specific mutations in *gag,* suggesting that the chemotype does not directly target Gag. We hypothesize that this chemotype may represent a first-in-class inhibitor of virus production that acts by targeting a viral-host complex important for HIV-1 Gag assembly.

## INTRODUCTION

In an era in which HIV-1 vaccine trial results have been disappointing, antiretroviral (ARV) drugs stand out as a stunning success story. Treatment with combined antiretroviral therapy (ART) over the past three decades has resulted in enormous reductions in morbidity and mortality from HIV-1 infection, with 62% of the 38 million people living with HIV receiving antiretroviral drugs in 2018 (1). ART is also the mainstay of PREP, a remarkably successful approach to preventing HIV-1 infection (1, 2). Additionally, ART will be important for future HIV cure strategies. Unfortunately, the success of HIV ART is at risk of being undermined by the increasingly serious problem of HIV-1 drug resistance (3). Virologic failure is seen in up to 20% of individuals receiving first-line ART in low and middle-income countries (4), with up to half of first-line ART failures in sub-Saharan Africa involving resistance to all three drugs in tenofovir-containing regimens (5). Moreover, the prevalence of drug-resistant HIV-1 is predicted to increase substantially over time as more people receive treatment (6). Indeed, experts have drawn analogies between the future of ART and the current crisis of multidrug resistant tuberculosis (7). For this reason, identifying novel targets in the HIV-1 life cycle and candidate small molecules that inhibit these targets is a high priority, with the goal of driving development of new ARV drugs.

Identification of new druggable targets in the poorly understood stages of the viral life cycle will be particularly important for development of drugs aimed at HIV-1 strains resistant to currently available drugs. Mapping the targets of drugs currently in use onto the HIV-1 life cycle reveals a striking lack of drugs that target intracellular events of assembly (8). Currently available ARV drugs target the maturation, entry, and post-entry stages of the viral life cycle. In contrast, no approved ARV drugs target intracellular viral late events (purple bar in Fig. 1A), which include post-translational events of immature capsid assembly. Formation of a single infectious virus requires assembly of ∼3000 copies of the HIV-1 Gag protein, which are synthesized and oligomerize in the cytoplasm and subsequently target to the plasma membrane, where they encapsidate the HIV-1 genome while multimerizing to form the immature capsid. The immature capsid then undergoes budding, release, and maturation (Fig. 1A). Interest in targeting HIV-1 assembly has been high, leading to numerous drug screens in recent years (9, 10). Of the compounds identified in these screens, the most successful have been those that bind to the capsid protein (CA). CA is critical at multiple stages of the viral life cycle including immature capsid assembly, maturation, infectivity, and post-entry events. Interestingly, potent small molecules that target CA do not selectively inhibit immature capsid assembly (reviewed in (9)) – instead they either act preferentially on the mature capsid, targeting virion infectivity and post-entry events (e.g. PF74 (11–16)), or act broadly on Gag synthesis, assembly, and post-entry events (e.g. GS-CA1 (17)) (Fig. 1A). Indeed, to our knowledge, there are no reports of a potent small molecule identified in any screen that selectively inhibits intracellular HIV-1 late events.

**FIG 1.**
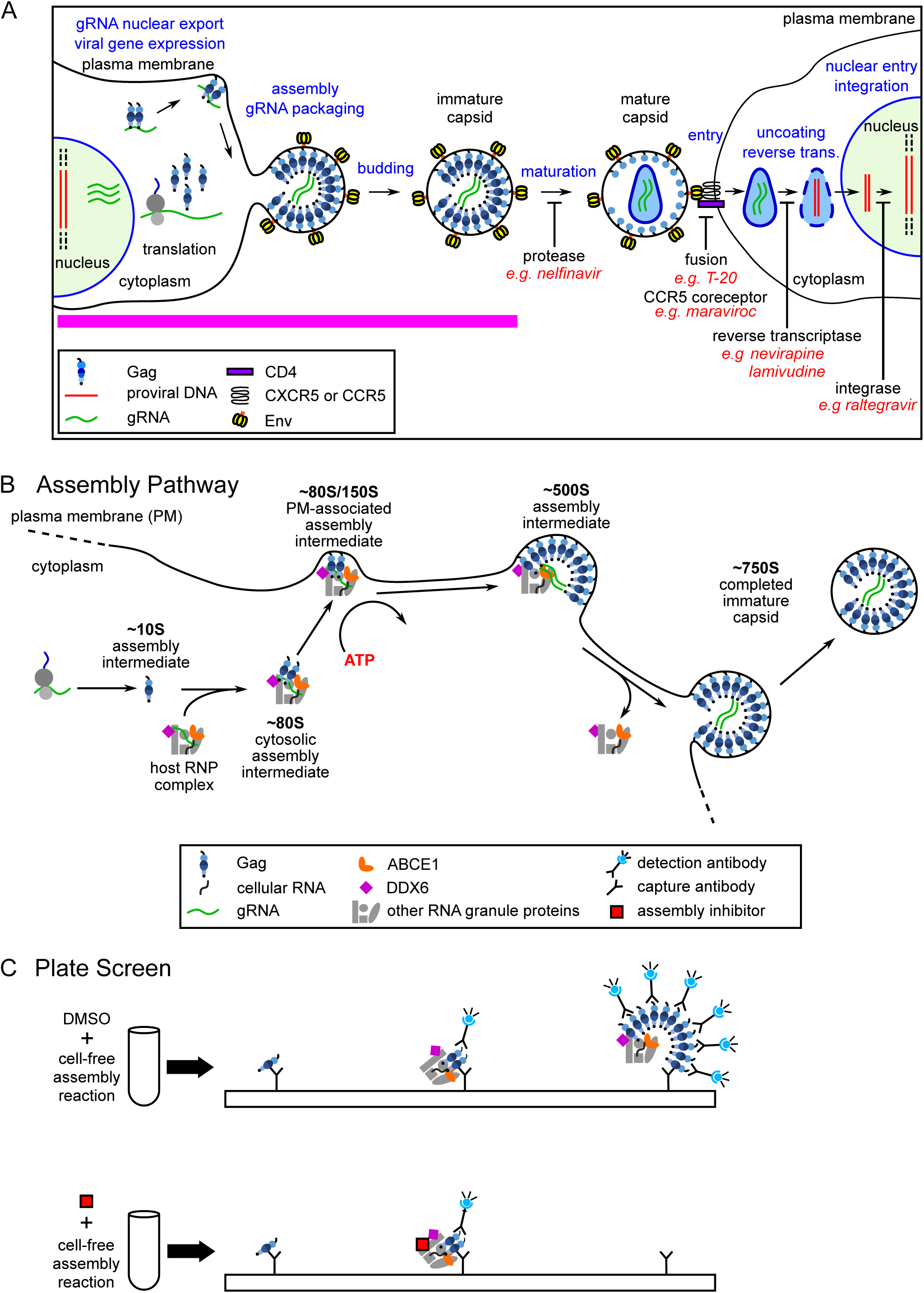
The HIV-1 life cycle, assembly pathway, and assembly plate screen. (A) Schematic showing the HIV-1 lifecycle, with expression of the integrated provirus, followed by the late and early phases of the lifecycle, ending with integration in a newly infected cell. The different stages of the virus lifecycle are indicated in blue. Examples of ARV drugs currently in use are in red, with black labels under blockade arrows indicating the targets of these drugs. Pink line indicates late events in the viral lifecycle that are not targeted by currently approved ARV drugs. (B) Schematic showing the host-catalyzed HIV-1 assembly pathway, starting with Gag synthesis and formation of the early ∼10S assembly intermediate. Next, the cytosolic ∼80S intermediate forms when ∼10S Gag co-opts a host RNP complex that contains ABCE1 and the RNA granule protein DDX6, two host enzymes that have been shown to facilitate assembly. The ∼80S assembly intermediate appears to target to the plasma membrane where Gag multimerization continues resulting in formation of the ∼150S and subsequently the ∼500S late assembly intermediate. When assembly of ∼750S immature capsid is completed, the host RNP complex is released. References are in the text. (C) Schematic showing the cell-free protein synthesis and assembly plate screen that was utilized to identify small molecule inhibitors of the host-catalyzed pathway of HIV-1 assembly. Briefly, anti-Gag antibody (capture antibody) binds Gag monomers, oligomers, and multimers generated in a cell-free assembly reaction. The same Gag antibody is used as a detection antibody that binds to captured oligomers and multimers, but not monomers, in proportion to the amount of multimerization, thereby generating a larger fluorescent signal when multimerization occurs. The upper diagram shows anti-Gag antibodies capturing and detecting Gag oligomers and multimers in assembly intermediates formed during an HIV-1 cell-free assembly reaction carried out in the presence of DMSO, which does not inhibit assembly. The lower diagram shows that adding an assembly inhibitor at the start of the cell-free reaction causes fewer Gag oligomers and multimers to be produced, thereby reducing the detection antibody signal relative to signal in the DMSO control.

We hypothesized that one reason for the failure to identify potent and selective inhibitors of HIV-1 assembly might be that previous screens were either very broad or very narrow – i.e. they encompassed the entire viral life cycle (e.g.(12, 18)) or they were based solely on multimerization of recombinant CA-derived peptides *in vitro* (e.g.(19–22)). We further hypothesized that a screen that focuses only on events of Gag assembly but includes known cellular facilitators of immature HIV-1 capsid assembly could be more successful than other screens in identifying a potent and selective inhibitor of intracellular events in HIV-1 assembly. Specifically, while recombinant Gag is able to assemble into immature capsid-like particles in the absence of host proteins (reviewed in (23)), two decades of studies support a different model for HIV-1 assembly in cells – one in which Gag assembles into immature capsids via a pathway of assembly intermediates containing viral proteins as well as host proteins that act catalytically to promote HIV-1 capsid assembly (e.g. (24–34); Fig. 1B). This model suggests that to succeed in the hostile environment of the cytoplasm, Gag may have evolved to utilize host proteins to catalyze Gag multimerization, promote RNA packaging, and sequester assembly within host complexes where nascent virions would be less vulnerable to host defenses. If this host-catalyzed model of capsid assembly in the cytoplasm is valid, then a screen that recapitulates this pathway might succeed in identifying new druggable targets.

Indeed, a precedent exists for a screen that recapitulates a host-catalyzed assembly pathway enabling identification of a novel antiviral compound and target. Previously our group, in collaboration with investigators at the Centers for Disease Control and Prevention, used a cell-extract-based screen that recapitulated an intracellular assembly pathway for rabies virus (RABV) to identify the first reported small molecule inhibitor of RABV replication in cell culture (35). Notably, this small molecule binds to a multiprotein complex that contains ATP-binding cassette protein E1 (ABCE1), a host enzyme identified in HIV-1 assembly intermediates, suggesting that similar host complexes may be involved in the assembly of diverse viruses.

Motivated by the RABV precedent, we developed cell-extract-based assembly pathway screens for other viruses and used them to identify a novel and potent antiretroviral chemotype that inhibits HIV-1 replication, represented by the small molecule PAV-117 and its more potent analog PAV-206. Here we present evidence suggesting that this chemotype inhibits HIV-1 replication by interfering with HIV-1 assembly. We found that a tagged analog of these compounds colocalizes with Gag *in situ*, suggesting that they may target HIV-1 Gag. However, surprisingly, compound-specific mutations did not arise in *gag* or *pol* after 37 weeks of PAV-206 selection in HIV-1-infected human cells, in stark contrast to nelfinavir selection examined in parallel, arguing against Gag being the direct target of PAV-206. Additional imaging studies shed more light on our failure to identify viral resistance mutations by demonstrating that the tagged PAV-206 analog also colocalizes with two host proteins associated with Gag in intracellular HIV-1 capsid assembly intermediates, the enzymes ABCE1 and DEAD-box helicase 6 (DDX6). While the exact target and mechanism of action of PAV-117/PAV-206 remain unclear, these studies raise the possibility that they represent a class of novel host-targeting ARV small molecules that inhibits HIV-1 virus production by acting on host-protein-containing capsid assembly intermediates.

## RESULTS

### Development of a novel screen for HIV-1 assembly inhibitors based on previous studies of the intracellular HIV-1 capsid assembly pathway

Our interest in using the cell-free assembly system as a drug screen stems from an early study that utilized this system to generate the first evidence that immature HIV-1 capsids assemble via a host-catalyzed pathway of intermediates (28). Adapted from the *in vitro* protein synthesis systems that were used to identify signal sequences (36), the cell-free HIV-1 assembly system supports *de novo* synthesis of HIV-1 Gag polypeptides from a Gag mRNA using energy substrates, amino acids, and a cellular extract that provides host factors required for Gag translation as well as post-translational events of Gag assembly. When programmed with wild-type Gag mRNA, this system produces particles that closely resemble completed immature HIV-1 capsids produced by provirus-expressing cells, judging by their ultrastructural appearance as well as in their size and shape (as defined by a sedimentation value of ∼750S; (28)). Pulse-chase studies in cell-free assembly reactions revealed sequential progression of Gag through complexes of increasing size (∼10S to ∼80S/150S to ∼500S to ∼750S), consistent with these smaller complexes being intermediates in the formation of the ∼750S completely assembled immature capsid. Moreover, Gag mutants that had been defined by others to be assembly-defective in cells are arrested at specific steps of the cell-free assembly pathway, with assembly-competent Gag mutants progressing through the entire pathway (28, 37). Notably, biochemical analysis demonstrated that post-translational events in this assembly pathway required ATP, indicating that HIV-1 immature capsid assembly in cells is energy-dependent (28).

While initially identified in a cell-free system, the HIV-1 capsid assembly pathway has been largely studied in cellular systems in the last two decades. Key features of the assembly pathway were validated in cells expressing the HIV-1 provirus (reviewed in (32)), including the sequential progression of Gag through the pathway of assembly intermediates (26, 32), the energy dependence of the pathway (25), and the arrest of known assembly-defective Gag mutants at specific steps in the pathway (25-28, 32, 33, 38). The energy dependence of immature capsid assembly, which has been confirmed by other groups (39), was subsequently explained by the finding that the assembly intermediates contain at least two host enzymes that facilitate assembly - the ATPase ABCE1 and the RNA helicase DDX6 (30, 34). Other studies suggest that packaging of the HIV-1 genome appears to occur in the assembly intermediates (24, 32), and that other lentiviruses utilize analogous assembly pathways (25, 31). Immunoprecipitation and imaging studies confirmed the association of ABCE1 and DDX6 with assembling Gag and identified two other enzymes in assembly intermediates, AGO2 and DCP2 (24, 26, 27, 30–33). The finding that assembly intermediates contain proteins typically found in host RNA granules, e.g. DDX6, AGO2, and DCP2, supports a model in which assembling Gag co-opts a unique subclass of host ribonucleoprotein (RNP) complexes that are related to RNA granules but differ from well-studied RNA granules such as P bodies and stress granules. RNA granules are a diverse group of ribonucleoprotein complexes involved in all aspects of host RNA metabolism except protein synthesis. Sequestration of immature HIV-1 capsid assembly events in such host RNP complexes could allow HIV-1 assembly to be protected from the innate immune system while at the same time allowing assembling Gag access to the viral genome, which is found in these complexes (24, 32), and to utilize enzymes that could facilitate packaging and assembly (Fig. 1B).

While the cellular studies described above demonstrated the physiological relevance of the HIV-1 assembly pathway, we reasoned that the cell-free system in which it was identified could hold promise as a drug screen. A cell-free assembly drug screen would have the advantage of recapitulating evolutionarily conserved viral-host interactions critical for assembly while being much less expensive than drug screens that utilize infected cell lines in culture. Additionally, the cell-free system can be adapted for a moderately high-throughput format. Together, these features of a cell-free assembly pathway drug screen allow a large number of compounds to be screened in a cost-effective manner.

For these reasons, we adapted the HIV-1 cell-free assembly reaction to generate a drug screen (Fig. 1C) similar to the RABV drug screen (35). Cell free reactions, upon completion, contain assembly intermediates and fully assembled immature capsid-like particles, unless performed in the presence of an inhibitor of assembly, in which case Gag will be largely unassembled (i.e. monomers and dimers). Plate-bound anti-Gag antibodies capture monomeric, dimeric, and multimerized Gag from the reaction. To determine how much Gag multimerization occurred during the reaction, a soluble anti-Gag detection antibody is added. The detection antibody does not bind Gag monomers (since the monomer antigenic site is already bound by capture antibody) but would be expected to bind to unoccupied Gag proteins in multimers. Signal from bound detection antibody is quantified using a fluorescent detection system, with more signal being generated by Gag multimers. Thus, an effective inhibitor of Gag multimerization will be recognized by a reduction in detection antibody signal (Fig. 1C). Because we are interested in inhibitors of assembly rather than protein synthesis, we utilized a counter screen to eliminate from further consideration all small molecules that inhibit translation of GFP in cell-free reactions.

### Identification of a potent inhibitor of HIV-1 replication

Once this cell-free HIV-1 assembly screen and counter screen were successfully established and validated, similar cell-free screens were established for quantifying assembly of the capsids of seven other viruses (Fig. 2A). A compound library of ∼150,000 small molecules with drug-like characteristics was screened against these eight cell-free assembly assays, resulting in a master hit collection of 249 small molecules that inhibited assembly in one or more virus assembly screen(s), but did not significantly inhibit GFP translation.

**FIG. 2.**
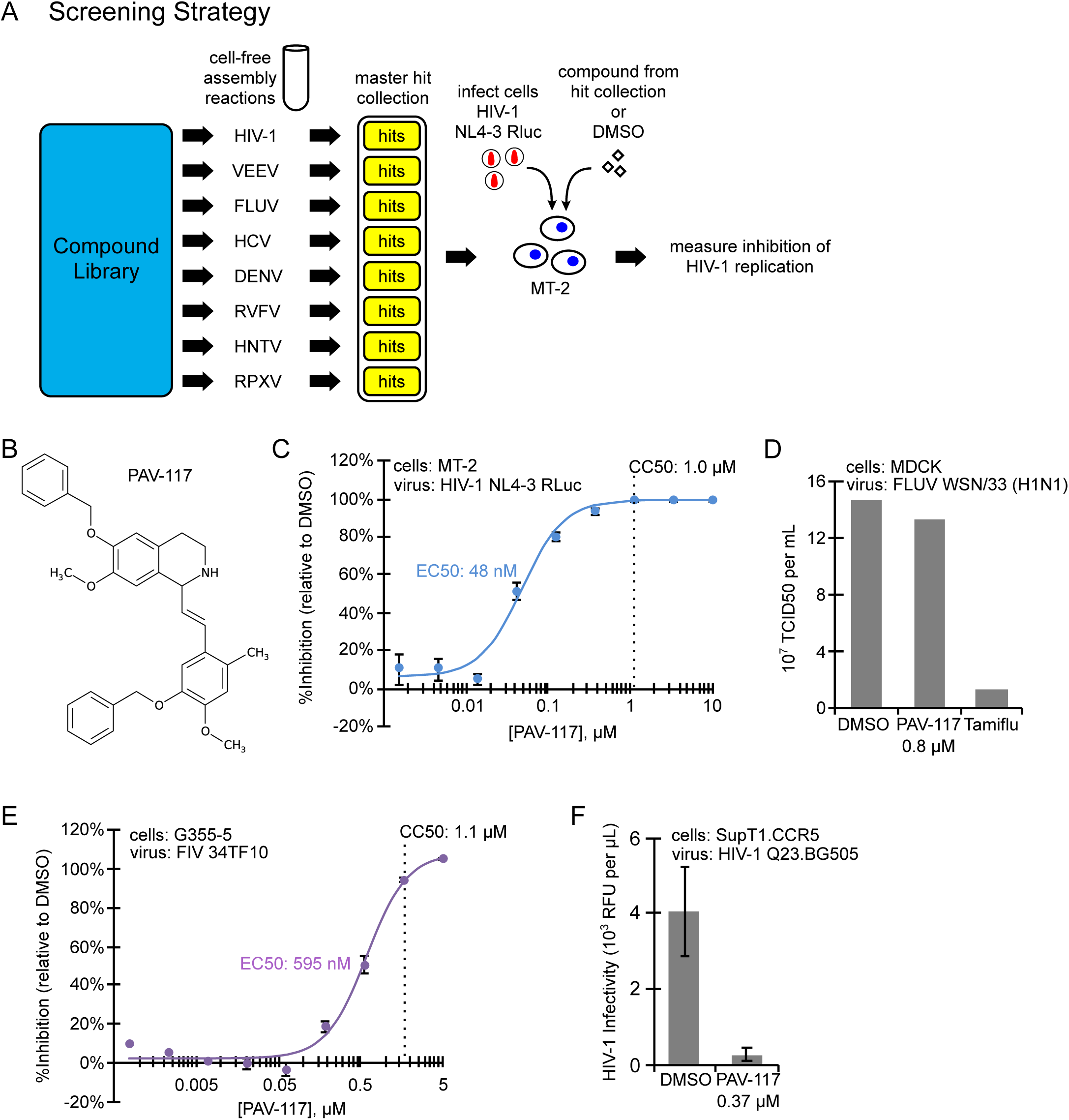
PAV-117, identified using cell-extract-based assembly screens, inhibits replication of HIV-1, but not FIV or FLUV, in cell culture. (A) Schematic showing the cell-free assembly screening strategy used to identify PAV-117. Small molecules from a library of 150,000 compounds were assayed in cell-free screens that recapitulate assembly of eight different viral capsids, with each screen being analogous to the HIV-1 screen in (Fig 1C). Results from the library screens led to generation of a master hit collection of 249 small molecules that displayed inhibitory activity in one or more of the eight cell-free assembly screens. Compounds in the master hit collection were assayed for inhibition of HIV-1 replication in MT-2 T cells, with PAV-117 identified as the compound with optimal EC50, CC50, and other characteristics in the MT-2 assay. (B) The chemical structure of PAV-117, a tetrahydroisoquinolone. (C) To determine the EC50 of PAV-117 against HIV-1 replication, a dose-response curve for inhibition of HIV-1 replication by PAV-117 was generated by treating human MT-2 T cells with indicated doses of PAV-117 followed by infection with a replication-competent HIV-1 NL4-3 RLuc reporter virus (MOI of 0.02). After 96 h of spreading infection, luciferase activity was measured as an indicator of HIV-1 replication and is displayed as inhibition of replication relative to DMSO-treated controls (% inhibition). CC50 was determined in parallel using uninfected MT-2 T cells and is marked by a vertical dashed line. Error bars show standard error of the mean (SEM) determined from three replicates. (D) The graph shows quantification of infectious FLUV in MDCK cells treated with DMSO or with either PAV-117 or Tamiflu at a concentration 20-fold higher than the EC50 for each drug (0.8 μM for PAV-117, 10 μM for Tamiflu). Treated cells were infected with FLUV (strain WSN/33(H1N1), MOI of 0.001) and viral titer was measured after 24 h by TCID50. (E) To determine the EC50 of PAV-117 against FIV replication, a dose-response curve for inhibition of FIV replication by PAV-117 was generated by treatment of G355-5 cells with indicated doses of PAV-117 followed by infection with FIV 34TF10 (1954 nU of RT activity per well). After 144 h of spreading infection, RT activity was measured and used to calculate inhibition of FIV replication relative to DMSO controls (% inhibition). CC50 was determined in parallel using uninfected G355-5 cells and is marked by a vertical dashed line. Error bars show SEM determined from three replicates. (E) Quantification of replication of an HIV-1 primary isolate. SupT1.CCR5 cells were treated with 0.37 μM PAV-117 or DMSO followed by infection with HIV-1 Q23.BG505, a CCR5-tropic subtype A molecular clone. HIV-1 replication proceeded for 96 h followed by measurement of HIV-1 infectivity in the culture supernatant using the MUG assay in TZM-bl cells.

Compounds in the master hit collection were further screened for their ability to inhibit replication of HIV-1 in the MT-2 human T cell line. The compound with the most favorable characteristics in this assay was PAV-117, a tetrahydroisoquinolone derivative with excellent drug-like properties (Fig. 2B). PAV-117 inhibits replication of HIV-1 NL4-3 in MT-2 cells with a 50% reduction in virus replication (EC50) of 48 nM and a 50% reduction in cell viability (CC50) of 1.0 μM (Fig. 2C). The nanomolar EC50 and a selectivity index (CC50/EC50) of 21 make PAV-117 an excellent compound for further optimization. Notably, PAV-117 was not active against influenza A virus WSN/33 [H1N1] (FLUV) suggesting that it is not an antiviral that acts broadly on all enveloped viruses (Fig. 2D). Moreover, PAV-117 was also relatively inactive against feline immunodeficiency virus (FIV), a non-primate lentivirus that is related to HIV-1, indicating specificity even among lentiviruses (Fig. 2E). PAV-117 was active against a primary isolate of HIV-1 (Fig. 2F), indicating that its antiretroviral activity is not restricted to laboratory isolates of HIV-1.

### Defining the step in the viral life cycle at which the novel small molecule acts

Since other types of assembly inhibitor screens had identified compounds that act on viral early events as described above, we next asked whether PAV-117 inhibits HIV-1 replication by acting on entry or post-entry events in the HIV-1 life cycle. For this assay, we infected MT-2 cells, in the presence of PAV-117 or DMSO, with a luciferase-encoding HIV-1 reporter virus (HIV-1 pNL4-3 RLuc Δ*env*) that is pseudotyped with HIV-1 Env and undergoes only one round of replication (Fig. 3A, diagram). At non-toxic concentrations, PAV-117 did not reduce luciferase activity in this assay (Fig. 3A, graph; EC50 of 930 nM; CC50 of 1.1 μM), indicating that it does not act on early events in the HIV-1 life cycle. The effect of PAV-117 on viral late events (events that occur after integration) was assayed using chronically infected H9 T cells, which produce infectious virus from integrated HIV-1 provirus (Fig. 3B, diagram). In this assay, PAV-117 inhibited production of infectious virus in the culture supernatant, with an EC50 of 104 nM (Fig. 3B, black dose-response curve in graph). Thus, PAV-117 activity can be entirely attributed to inhibition of viral late events. To determine whether PAV-117 results in production of virus-like particles (VLP) that were non-infectious, we quantified production of p24 Gag in VLP pelleted from cell culture supernatants of chronically infected H9 cells treated with PAV-117 or DMSO. Finding that PAV-117 reduces virus infectivity without altering the amount of p24 Gag in virus pellets would indicate that PAV-117 results in production of non-infectious 10 VLP. However, what we actually observed was the opposite - that inhibition of p24 production was similar to inhibition of supernatant infectivity (EC50 of 113 nM vs. 104 nM, respectively; compare blue vs. black curve in Fig. 3B graph), indicating that the compound blocks virus production. Western blots of VLP in supernatants also showed no obvious change in the ratio of p55 to p24 with increasing PAV-117 concentration relative to DMSO treatment (Fig. 3B, blot), arguing that it does not act on Gag cleavage required for virus maturation.

**FIG. 3.**
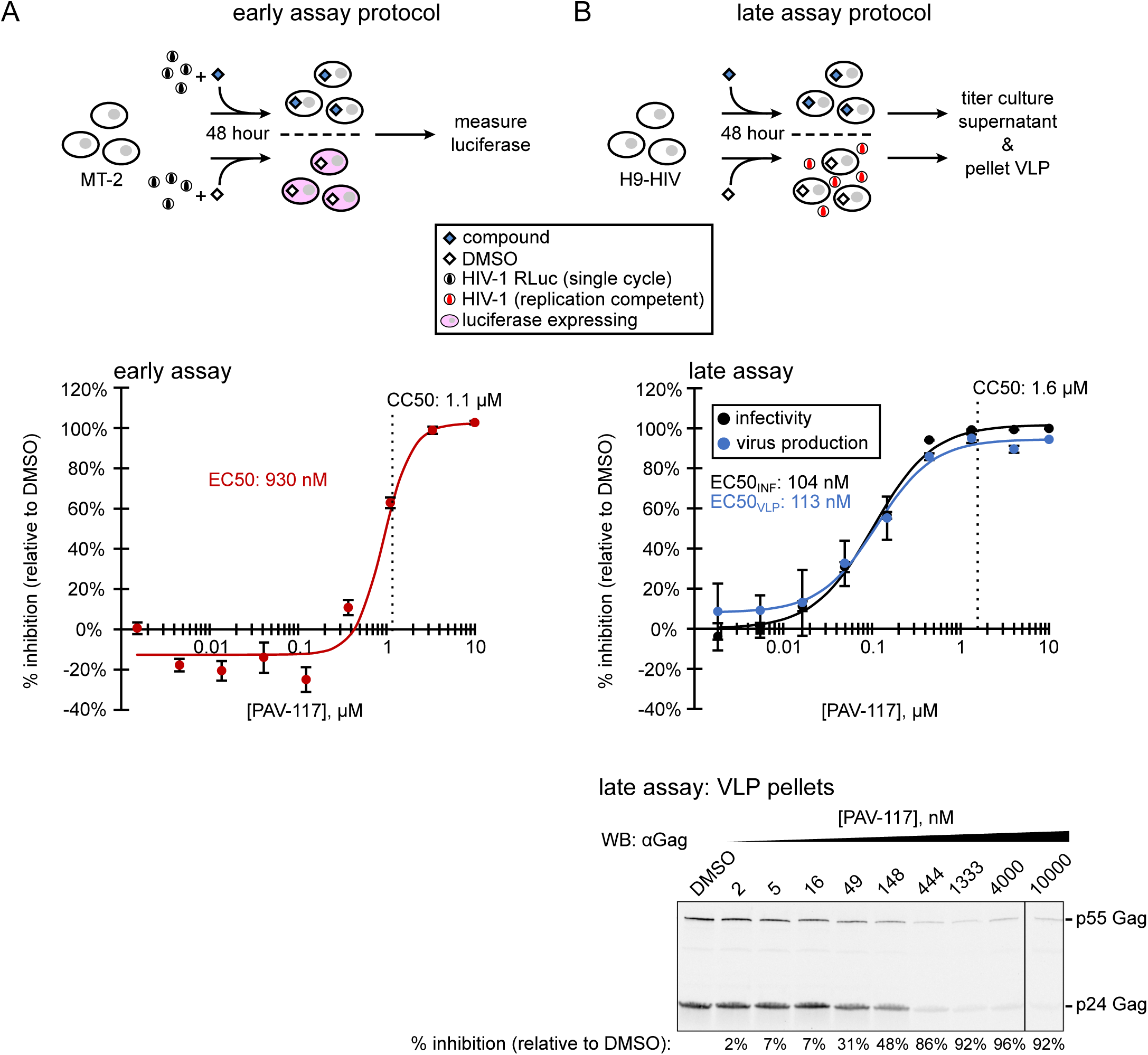
PAV-117 acts late in the HIV-1 life cycle, inhibiting virus production but not specific infectivity. (A) Schematic of the early assay, which measures effects on viral entry through early viral gene expression. MT-2 cells were infected with the single-round HIV-1 pNL4-3 RLuc virus (*env*-deleted and pseudotyped with HIV-1 NL4-3 Env) in the presence of compound or DMSO. After 48 h, luciferase activity was measured and used to calculate inhibition of HIV-1 replication relative to the DMSO control (% inhibition). The graph shows the dose-response curve for inhibition of HIV-1 early events by PAV-117 that was generated using this assay and used to determine the EC50. CC50 was determined in parallel using uninfected MT-2 T cells and is marked by a vertical dashed line. Error bars in graph show SEM determined from three replicates. (B) Schematic of the late assays, which measure effects on viral late events starting with expression of Gag and GagPol through virus release and maturation. Chronically infected H9 T cells (H9-HIV) were treated with either compound or DMSO, and media collected 48 h later were used for two assays: 1) to quantify inhibition of HIV-1 infectivity relative to DMSO control by titering on TZM-bl cells (black curve, used to calculate EC50 for inhibition of infectivity) and 2) to quantify inhibition of virus production by pelleting virus for western blot (WB) with antibody to HIV-1 Gag (αGag; blue curve, used to calculate EC50 for inhibition of virus production). The CC50 was determined in the inhibition assay and is marked by a vertical dashed line. A representative αGag WB of virus pellets is shown below the dose-response graph, with DMSO treatment or concentration of PAV-117 indicated above each WB lane and % inhibition of virus production (relative to the DMSO-treated control) indicated below each lane. Error bars in (B) and (C) show SEM determined from two replicates.

The finding that the entire effect of PAV-117 is due to inhibition of virus production (Fig. 3) indicated that PAV-117 likely acts on an intracellular viral late event. Such events include transcription of the full-length HIV-1 mRNA, translation of this mRNA to produce Gag and GagPol, assembly of Gag to produce immature capsids, and budding at the plasma membrane to release immature virus. We hypothesized that PAV-117 likely acts on post-translational events in assembly of immature HIV-1 capsids for two reasons: first, because a counter screen had been utilized to eliminate inhibitors of translation; and second, because the cell-free screens that identified PAV-117 recapitulate capsid protein assembly but not virus budding and release. To test this hypothesis, we analyzed acutely infected MT-2 cells that had been treated for 48 h with DMSO or PAV-117 at 0.25 nM vs. 0.75 nM (∼EC70 and ∼EC90). As expected, we observed a dose-dependent reduction in p24 release into the supernatant (Fig. 4A). Consistent with PAV-117 not being an inhibitor of translation, steady-state levels of actin and GAPDH were minimally affected (Fig. 4B). At the highest PAV-117 concentration tested, some reduction in steady-state Gag levels was observed (Fig. 4A), but it was much less than the reduction in VLP production. To further define where PAV-117 acts, we also examined steady-state levels of HIV-1 capsid assembly intermediates, whose distinct S values allow them to be separated by velocity sedimentation. PAV-117 had little to no effect on steady-state levels, as % of total Gag, of the ∼10S and ∼80S/150S intermediates but resulted in a dose-dependent decrease in the steady-state level of the ∼500S intermediate (Fig. 4C). While the steady-state data in Fig. 4 provide a snapshot of intracellular Gag levels at only one point in time, taken together with the dramatic reduction in cumulative VLP production (across 48 h) in Fig. 3B, these data suggest that PAV-117 acts during virus assembly, most likely by inhibiting formation of the ∼500S assembly intermediate.

**FIG. 4.**
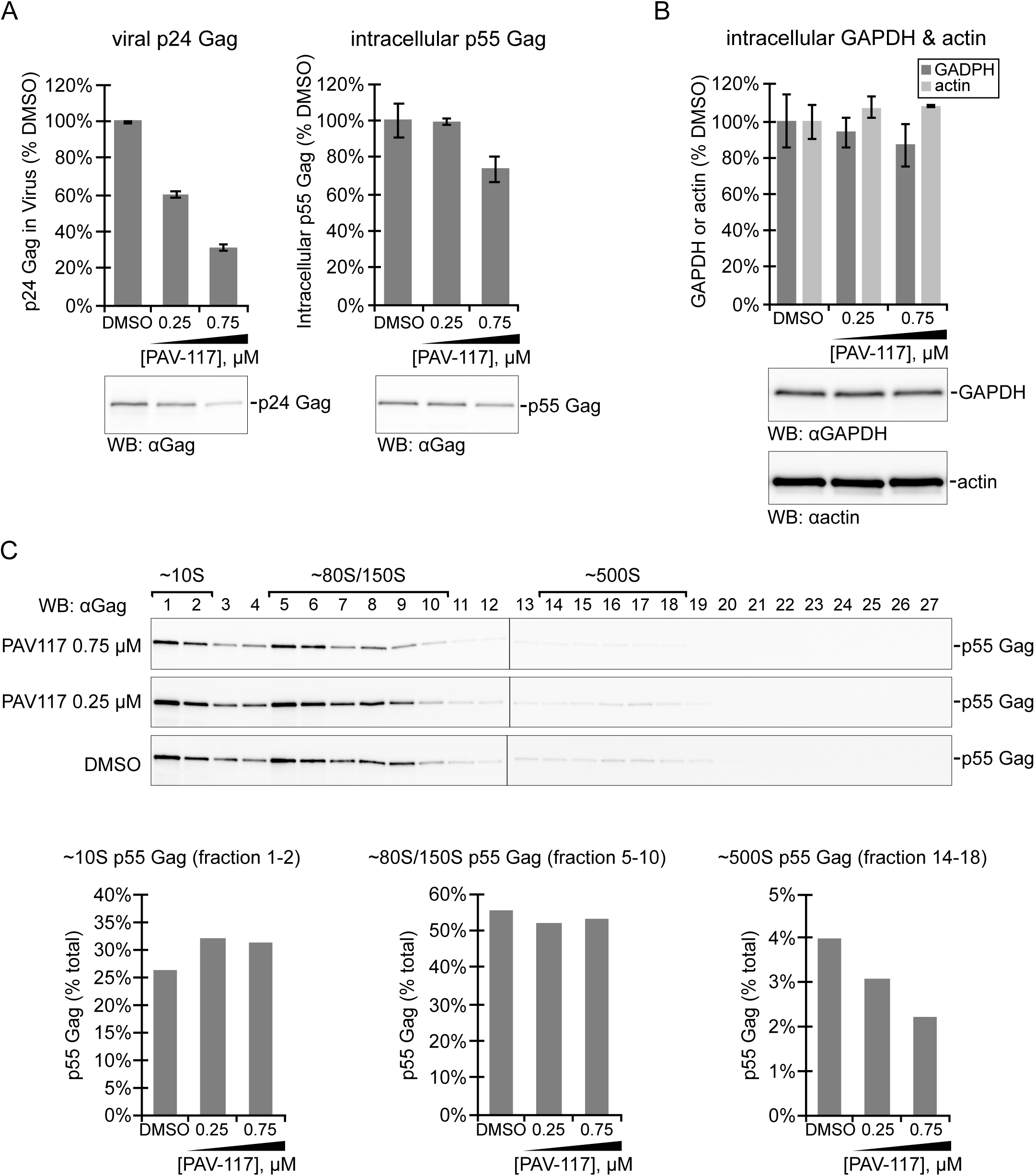
PAV-117 appears to act during the HIV-1 assembly pathway, after formation of the ∼80S intermediate. MT-2 cells were infected with HIV-1 LAI *pro*- Δ*env* (pseudotyped with HIV-1 NL4-3 Env) and treated with DMSO or the indicated concentrations of PAV-117 for 48 h. (A) Media and cell lysates were harvested to analyze effects on virus production and intracellular steady-state Gag levels, as indicated, using WB with Gag antibody (αGag) to quantify p24 or p55 Gag. (B) Cell lysates were also analyzed for intracellular steady-state levels of two cellular proteins, GAPDH and actin, by WB with αGAPDH and αactin. For (A) and (B), data in graphs are shown as % of DMSO-treated controls, with error bars showing SEM from two replicates, and representative WBs are shown below graphs, (C) To quantify intracellular steady-state levels of assembly intermediates, cell lysates from panels A and B were also analyzed by velocity sedimentation followed by WB of each gradient fraction with αGag. Fraction numbers are indicated above the WB panels, with migration of specific assembly intermediates indicated by brackets above. Graphs show quantification of p55 Gag in fractions containing the ∼10S, ∼80S/150S, and ∼500S intermediates as percent of total p55 Gag in the gradient. Expected migration of each protein in WB panels is indicated to the right.

### Generation of analogs of the novel small molecule inhibitor that are more potent or contain a tag

If PAV-117 acts during assembly, one might expect it to be associated with assembling HIV-1 Gag. To test if this is the case, we needed an analog of PAV-117 that contains a tag, such as biotin, to allow detection while retaining antiviral activity. To identify positions that could be used for a biotin tag, structure-activity relationships were analyzed with combinations of methyl, methoxy, or benzoyloxy substitutions in the R1, R2, and R3 positions of the pendant benzene ring (Fig. 5A). Interestingly, we observed that the presence of a methyl in the R1 position and a methoxy in the R2 position (PAV218 and PAV-206) resulted in reduced toxicity (higher CC50) and a higher selectivity index when analyzed in HIV-1 infected MT-2 T cells (Fig. 5A). PAV-206, which differs from PAV-117 only because it contains a methoxy rather than a benzoyloxy group in the R3 position, was particularly notable since it had a lower EC50 and higher CC50 than PAV-117, resulting in an excellent average SI of 61 in MT-2 cells (Fig. 5A, B) and 48 in HIV-1 infected PBMCs (Fig. 5B).

**FIG. 5.**
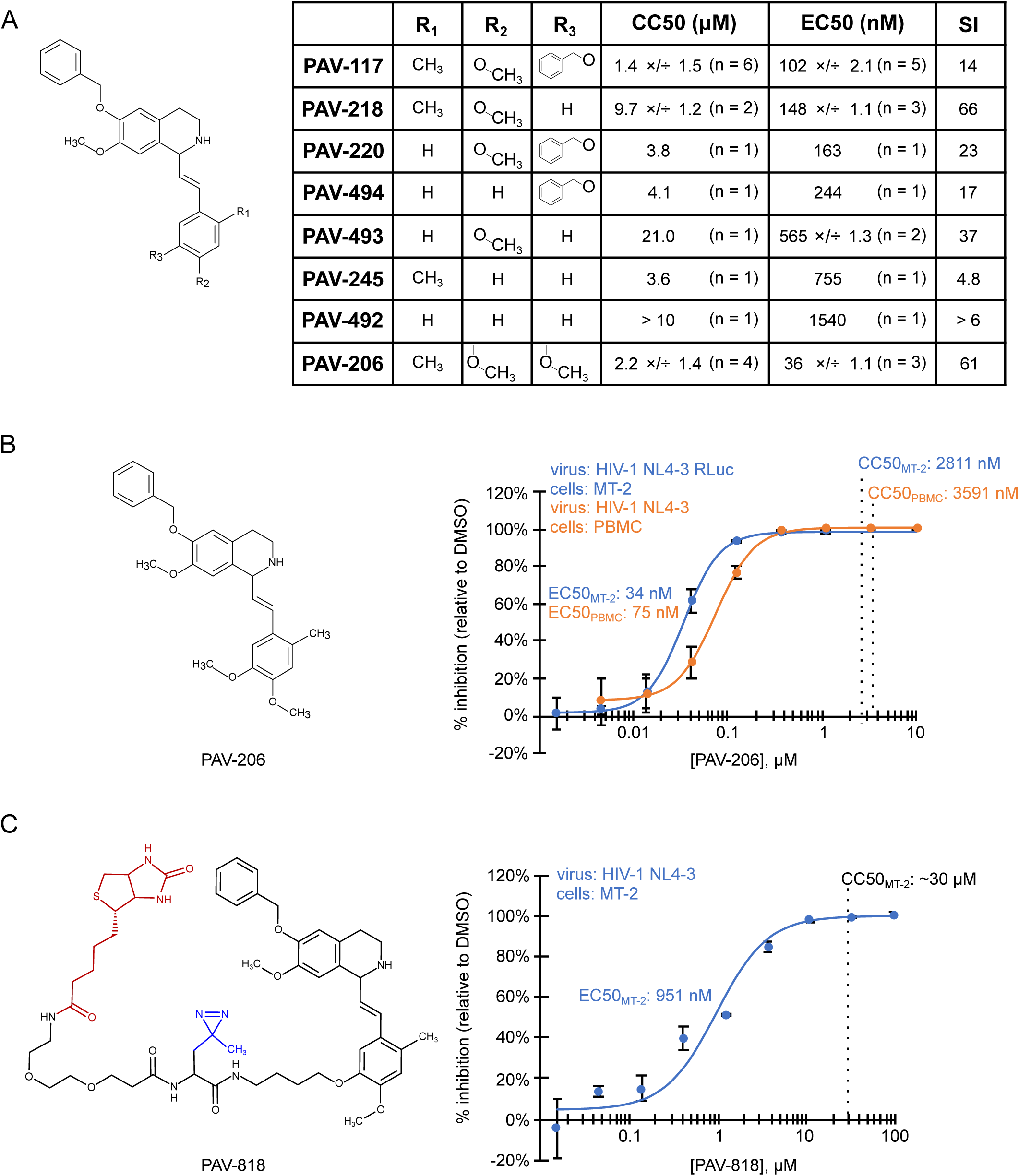
Analysis of structure-activity relationships identified an analog that potently inhibits HIV-1 replication in PBMCs and a site for tags. (A) The general chemical structure of PAV-117 analogs is shown at left indicating the R1, R2, and R3 positions in the pendant benzene ring. The table shows results obtained for analogs in which the R1, R2, and R3 positions contain hydrogen, methyl, methoxy, or benzoyloxy groups as indicated, including the EC50 for inhibition of HIV-1 replication in MT-2 cells and the CC50 in MT-2 cells (assays described in Fig. 2C). Values are shown as the average of multiple independent repeats ×/÷ GSD, with *n* = the number of independent repeats. Also shown is the selectivity index (SI), which is equivalent to CC50/EC50. (B) The structure of PAV-206 is shown at left. The blue dose-response curve shows inhibition of HIV-1 replication by PAV-206 in MT-2 T cells (using the assay described in Fig. 2C). The orange dose-response curve shows inhibition of HIV-1 replication by PAV-206 in PHA-activated PBMC infected with unmodified HIV-1 NL4-3 at an MOI of 0.008. (C) Shown on the left is the structure of PAV-818, the biotinylated analog of PAV-206, and on the right a dose-response curve for inhibition of HIV-1 replication by PAV-818 in MT-2 T cells (assays as in Fig. 2C). In the PAV-818 structure, the biotin moiety is shown in red. A diazirine group in the PAV-818 structure is shown in blue; this group was added for future crosslinking studies but is not used in the current study. For all graphs in (B) and (C), the indicated EC50s were determined from the dose-response curves; CC50s were determined in uninfected cells in parallel and are marked by vertical dashed lines. Error bars in (B) and (C) show SEM from three replicates.

In addition to identifying PAV-206, a more potent analog of PAV-117, the structure-activity relationship analysis led to a strategy for introducing a tag into PAV-117 (or PAV-206) by revealing that the R3 position can tolerate either a bulky group (as in PAV-117) or a hydrogen (as in PAV218) without significant loss of activity (Fig. 5A). For this reason, we introduced a biotin at this position, generating the PAV-818 analog (Fig. 5C), which is identical to PAV-117 and PAV-206 except at the R3 position. PAV-818 has a higher EC50 (951 nM) than the PAV-117 or PAV-206 analogs, but also has a higher CC50 (∼30 μM), generating an SI of ∼32 (Fig. 5C), which is slightly better than the SI of PAV-117. Thus, PAV-818 represents a biotinylated analog of PAV-206 and PAV-117 that can be utilized to visualize where these compounds localize within infected cells.

### A tagged analog of PAV-206 colocalizes with HIV-1 Gag *in situ* in infected cells

Next we utilized the biotinylated antiviral analog PAV-818 to ask if this family of compounds colocalizes with HIV-1 Gag as one would expect if they inhibit assembly, as suggested by data in Figs. 3 and 4. For this purpose, we used the proximity ligation assay (PLA), a technique which marks two antigens that are in close proximity with fluorescent spots. Briefly, in PLA, primary antibodies from two species are detected by species-specific secondary antibodies that are tagged with oligonucleotides. These oligonucleotides hybridize to connector oligonucleotides only if the antigens detected by the primary antibodies are within ∼40 nm of each other. Ligation of connector oligonucleotides and addition of a polymerase results in rolling circle amplification to generate a sequence that can be detected by a complementary oligonucleotide conjugated to a fluorescent probe (40). Thus, the fluorescent spots (in this case, red spots) represent sites where two antigens are colocalized. To determine whether PAV-818 colocalizes with HIV-1 Gag, we performed biotin-Gag PLA on 293T cells that were chronically infected with HIV-1 and treated with either the biotinylated antiretroviral PAV-818, a biotinylated compound that lacks antiviral activity (PAV-543; structure in Fig. S1), or the DMSO vehicle (Fig. 6A). Quantification of red fluorescent spots, which represent sites where biotinylated compound and Gag are colocalized, revealed 5.5-fold more spots in cells treated with 10 μM PAV-818 than in cells treated with an equivalent concentration of the control compound PAV-543, and a slightly greater difference relative to DMSO treated cells (Fig. 6B, C). Moreover, the number of biotin-Gag spots observed with PAV-818 treatment displayed a dose-response relationship (Fig. 6B, C). Controls revealed that only a background level of spots was observed when uninfected 293T cells were treated with PAV-818, as would be expected given that uninfected cells lack Gag (Fig. 6B, C). Additionally, as expected, when antibody to biotin was replaced with a nonimmune control antibody only a few spots were observed; similar results were observed when antibody to Gag was replaced with a nonimmune control (Fig. S2). Together these data provide evidence that PAV-818 colocalizes with Gag in HIV-1 infected human cells.

**FIG. 6.**
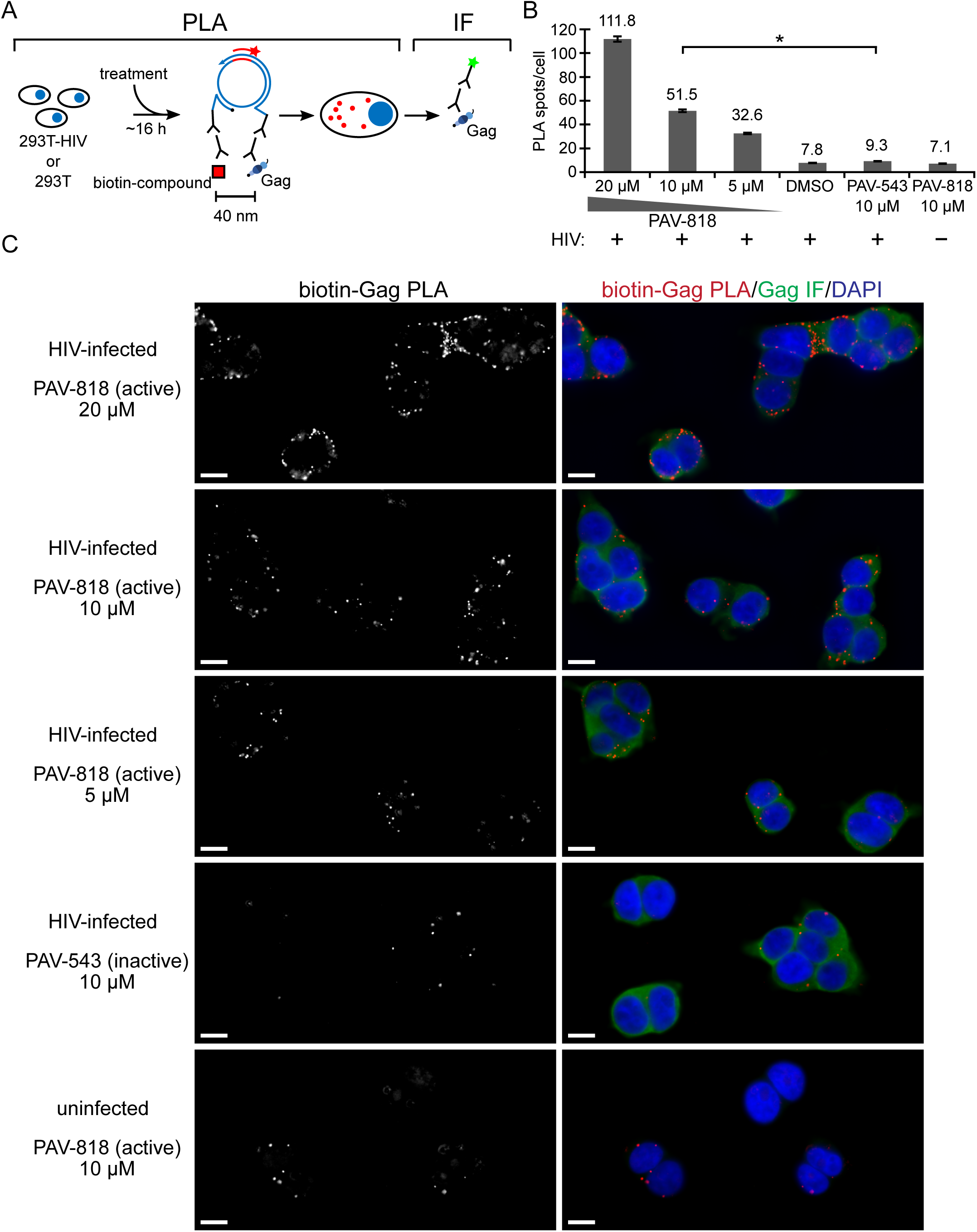
An antiviral analog of PAV-206 colocalizes with HIV-1 Gag *in situ*. (A) Schematic of the PLA approach for detecting colocalization of compound with Gag. 293T cells chronically infected with HIV-1 (293T-HIV) or uninfected 293T cells were treated with indicated amounts of PAV-818 (the biotinylated active compound), PAV-543 (the biotinylated inactive compound), or DMSO for 16 h. PLA was performed by incubating with primary antibodies (rabbit anti-biotin and mouse anti-Gag) followed by PLA secondary antibodies (anti-rabbit IgG coupled to (+) PLA oligo and anti-mouse IgG coupled to (-) PLA oligo). Addition of other PLA reagents leads to connector oligos linking the (+) and (-) oligos only if the primary antibodies are colocalized; this in turn results in the PLA amplification reaction. Addition of an oligo that recognizes a sequence in the amplified regions and is coupled to a red fluorophore (red star) results in intense spots at sites where biotinylated compound and Gag are colocalized *in situ*. Following PLA, indirect immunofluorescence (IF) is performed by adding secondary antibody conjugated to a green fluorophore (green star) to detect any unoccupied Gag antibody, thus marking Gag-expressing cells with low-level green fluorescence (B) The graph shows the average number of biotin-Gag PLA spots per cell for each condition, with + indicating HIV-1-infected cells and – indicating uninfected cells. Twenty fields were analyzed for each group (containing a total of 186 - 316 cells per group), with error bars showing SEM. * indicates a significant difference in the number of biotin-Gag PLA spots per cell when comparing treatment with PAV-818 vs. PAV-543, both at 10 μM (p < 0.001). (C) A representative field for each group quantified in (B) is shown, except for DMSO treatment. Fields on the left show biotin-Gag PLA spots alone in grayscale. To the right are the same fields shown as a merge of three color-channels: biotin-Gag PLA (red), Gag IF (green), and DAPI-stained nuclei (blue). Scale bars, 10 μm.

### Compound-specific resistance mutations were not observed in *gag* or *pol* after 37 weeks of selection with PAV-206

The colocalization of PAV-818 with HIV-1 Gag suggested a straightforward model in which Gag is the target of this assembly inhibitor. If this is the case, then selection for resistance to the compound should result in rapid acquisition of compound-specific mutations in the *gag* gene, analogous to what has been observed for all antiretrovirals that target viral gene products (41). To determine if development of resistance under selection could identify a viral target for this compound, we infected MT-2 T cells and passaged them under selection with PAV-206 for 37 weeks, increasing the drug concentration by two-fold whenever evidence of resistance was observed, and periodically sequencing virus to identify dominant mutations (Fig. 7A, sequencing events indicated by red arrows). We also selected with nelfinavir, a well-studied protease inhibitor, in parallel, as a positive control.

**FIG. 7.**
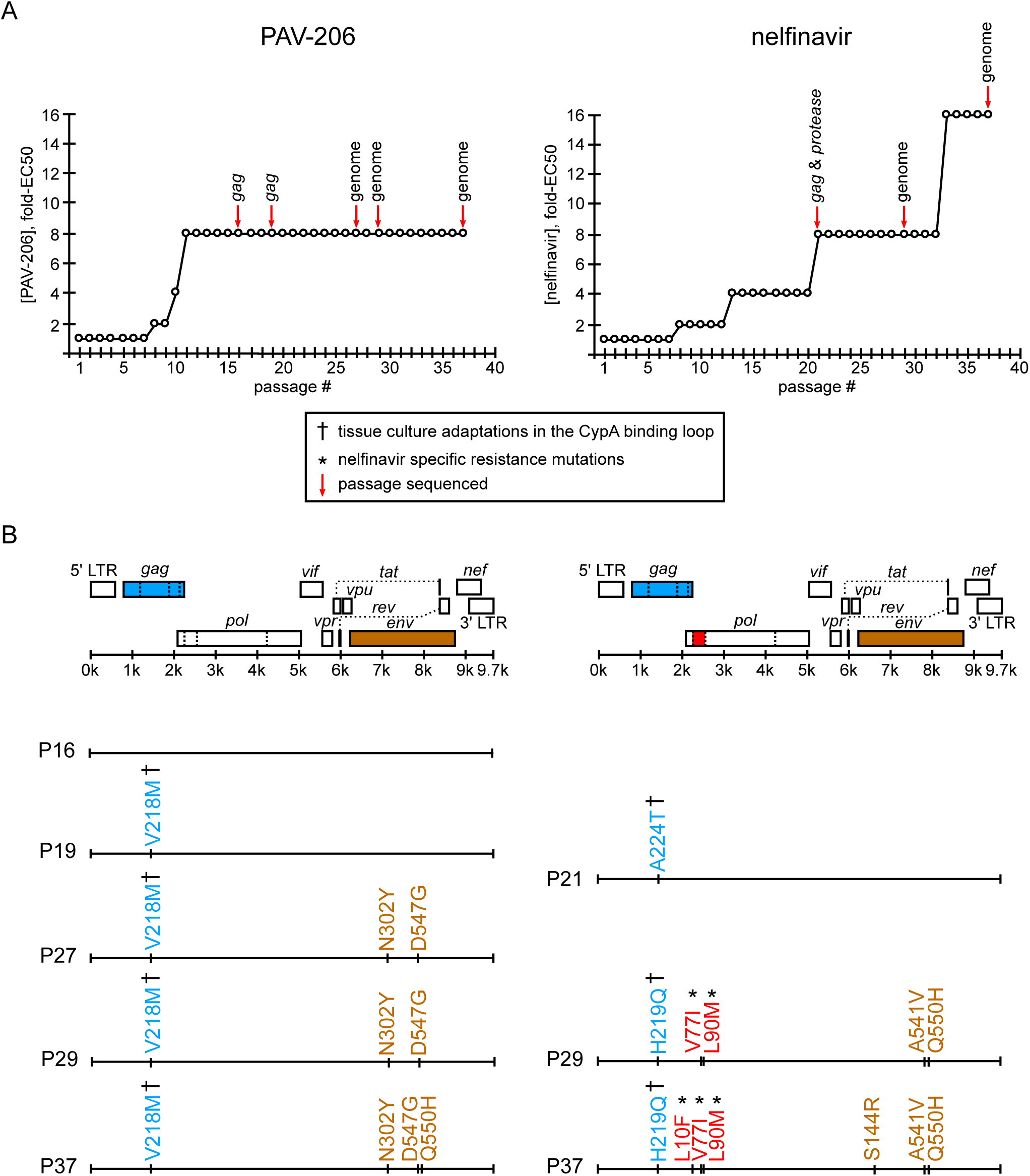
Resistance mutations in *gag* or *pol* were not observed upon selection with PAV-206 in cell culture. (A) A NL4-3 virus stock was passaged weekly for 37 weeks in the presence of either PAV-206 or nelfinavir starting at a concentration of 1 x EC50. The concentration of compound was increased two-fold when compound-treated cultures reached maximum viral cytopathic effect (CPE) in a timeframe similar to a parallel DMSO control. The graph shows the compound concentration as fold-EC50 vs. week of selection (passage #) for PAV-206 (left) or nelfinavir (right). Red arrows indicate passages where virus was amplified, either with sequencing of *gag* alone, *gag* & *protease*, or the full genome minus the LTRs (as indicated above red arrows). (B) For each selection (PAV-206 vs. nelfinavir), an HIV-1 genome map is shown, with positions of viral open reading frames (ORFs), 5’ LTR, and 3’ LTR, with a line below indicating their approximate nucleotide positions within the HIV-1 genome. In each case, ORFs in which dominant non-synonymous mutations emerged during selection are color-coded as follows: *gag*, blue; *pro*, red; *env*, brown. Beneath the HIV-1 genome maps, dominant non-synonymous amino acid mutations (identified through sequencing of passages indicated with red arrows in panel (A)) are shown according to their position in the genome map at the top of panel (B). The passage (P) in which these mutations were identified is indicated to the left of each genome map line. The amino acid mutations are color-coded according to the ORF in which they were found (colors 52 described above). As indicated in boxed legend, which applies to both panels, cyclophilin A binding loop mutations that have been previously identified as tissue culture adaptations are marked with a dagger symbol and previously described nelfinavir-specific mutations are marked with an asterisk (with all references for these in the text and Tables S1 and S2).

As would be expected, we observed that 37 passages in the presence of nelfinavir led to the emergence of three well known resistance mutations in HIV-1 protease: L90M, V77I, and L10F (indicated in red in Fig. 7B using HXB2 genome numbering; Table S1), with L90M being a primary resistance mutation and V77I and L10F being common secondary mutations that further boost nelfinavir resistance (42–46). L90M and V77I were first detected as dominant mutations at passage 29, just before the virus developed a high level of resistance, defined here as replication in a concentration of nelfinavir that is 16-fold higher than the EC50. L10F was detected later, becoming dominant at passage 37. In addition to these well described non-synonymous protease-resistance mutations, two non-synonymous substitutions in Gag emerged during nelfinavir selection (indicated in blue in Fig. 7B; Table S1). These substitutions, A224T and H219Q, are in the cyclophilin A binding loop of Gag and were detectable by passage 21 and 29, respectively. H219Q and other substitutions in the cyclophilin A binding loop (amino acids 217-225 in Gag) are polymorphisms in the Los Alamos Database (47, 48) that are known to modulate incorporation of cyclophilin A into virions when virus is passaged in cyclophilin-A-rich immortalized CD4+T cells even in the absence of drugs, thereby resulting in virion cyclophilin A levels that are optimal for replication in these cell lines (49, 50). From these data, we infer that in the nelfinavir selection group, a tissue-culture-adaptive cyclophilin A binding loop polymorphism initially emerged in Gag and likely led to improved fitness and replication in a drug concentration 8-fold greater than the EC50; this was followed later by nelfinavir-specific resistance mutations, which likely led to high level nelfinavir resistance.

Given the results with nelfinavir, we were surprised to find no dominant PAV-206-specific mutations in the *gag* or *pol* genes upon selection with PAV-206 in parallel. Notably, we did detect a dominant drug-independent, tissue-culture adaptive mutation in the cyclophilin A binding loop of Gag in the PAV206 selection, as had been observed with nelfinavir. Specifically, at passage 19, a V218M substitution in the cyclophilin A binding loop (Fig. 7A, B; Table S2). Like the cyclophilin A binding loop substitutions observed with nelfinavir selection, V218M improves viral fitness in cyclophilin-A-rich CD4+ T cells through optimizing cyclophilin binding (47). Unlike with nelfinavir selection, no other dominant non-synonymous mutations were observed in the *gag* or *pol* genes during 37 weeks of selection with PAV-206; moreover, high level resistance was also not observed (Fig. 7A, B), with an unsuccessful passage in a PAV-206 concentration that was 16-fold higher than the EC50 (J. Reed, unpublished observations).

While our primary interest was in dominant non-synonymous mutations in *gag* or *pol*, which include the most plausible targets of PAV-206, we also looked for mutations elsewhere in the HIV-1 genome. Interestingly, non-synonymous mutations in *env* were observed in both selection groups (A541V, Q550H, and S144R in the nelfinavir selection group, and N302Y, D547G, and Q550H in the PAV-206 selection group in Fig. 7A, B; Table S1, S2). These mutations are known to confer global replication advantage or serve as tissue culture adaptations. Specifically, A541V (observed at nelfinavir passage 29) was recently shown to confer broad escape from defects in virus replication caused by either virus mutations or antiretroviral drugs, most likely by increasing cell-to-cell transmission in T cell lines (51). Similarly, N302Y and D547G (observed at PAV-206 passage 27) are thought to increase fusion kinetics (52) and enhance fusogencity (53), respectively, and likely confer global replication advantages. Q550H, which arose during both PAV-206 and nelfinavir selection, has been observed in the absence of drug treatment (54) and is therefore likely to be a tissue culture adaptation. Thus, these *env* mutations are not specific to PAV-206 selection. Finally, we also observed three synonymous mutations arising in *env* upon PAV-206 selection (Table S2). Overall, we concluded from these resistance studies that replication in PAV-206 for 37 weeks failed to select for any PAV-206-specific resistance mutations in *gag* or *pol*, and therefore differed markedly from the nelfinavir selection control, which demonstrated the classic pattern of multiple resistance mutations emerging in the viral target of the drug. Consistent with this conclusion, we observed a high level of resistance in the nelfinavir selection group at passage 33 that correlated with emergence of nelfinavir-specific mutations in *pro*, while resistance to PAV-206 failed to rise above the moderate level that is likely due to tissue-culture adaptation and mutations that confer global replication advantage in T cells (Fig. 7A).

### PAV-206 colocalizes with host components of assembly intermediates, suggesting a host-targeting mechanism

Our studies to this point had shown that the PAV-206 chemotype colocalizes with Gag *in situ* (Fig. 6) and appears to inhibit Gag assembly (Fig. 4), but unexpectedly does not appear to target Gag or GagPol based on resistance studies (Fig. 7). This led us to hypothesize that PAV-206 and its analogs colocalize with host proteins as well as Gag, which would raise the possibility that PAV-206 targets a host protein or viral-host interface that is critical for assembly. This possibility is made more plausible by the fact that these compounds were identified through assembly screens that recapitulate host-catalyzed capsid assembly pathways. In the case of HIV-1 immature capsid assembly, two host proteins, ABCE1 and DDX6, are known to promote assembly and are associated with Gag, serving as markers of Gag-containing assembly intermediates (26, 30–34). Thus, we used PLA to next ask whether PAV-206 colocalizes with these Gag-associated host proteins.

293T cells chronically infected with HIV-1 were treated with either the biotinylated antiretroviral PAV-818, or the biotinylated inactive PAV-543, or DMSO and subjected to biotin-ABCE1 PLA (Fig. 8A). Quantification of red fluorescent spots representing sites where biotinylated compound and ABCE1 are colocalized *in situ*, revealed 14.3-fold more spots in cells treated with the antiviral PAV-818 (10 μM) than in cells treated with an equivalent concentration of PAV-543 and a similar difference relative to DMSO treatment (Fig. 8B, C). Additionally, the number of biotin-ABCE1 spots observed with PAV-818 treatment displayed a dose-response relationship (Fig. 8B, C). Interestingly, PAV-818 colocalized with ABCE1 as strongly in uninfected 293T cells as in HIV-1 infected human cells (Fig. 8B, C). As expected, controls in which antibody to biotin or to ABCE1 were individually replaced with a nonimmune control antibody generated very few PLA spots (Fig. S3). PAV-818 was also found to colocalize with DDX6 in chronically HIV-1 infected 293T cells by PLA, with 5.1-fold more spots observed in PAV-818-vs. DMSO-treated cells (Fig. 9A). As a control, we also showed that ABCE1 is colocalized with DDX6 in 293T cells chronically infected with HIV-1, as would be expected from our coimmunoprecipitation studies (30), using PLA with antibodies to DDX6 and ABCE1 (Fig. 9B). Thus, this novel antiretroviral compound colocalizes with HIV-1 Gag and with the host proteins ABCE1, and DDX6, all of which are components of HIV-1 assembly intermediates.

**FIG. 8.**
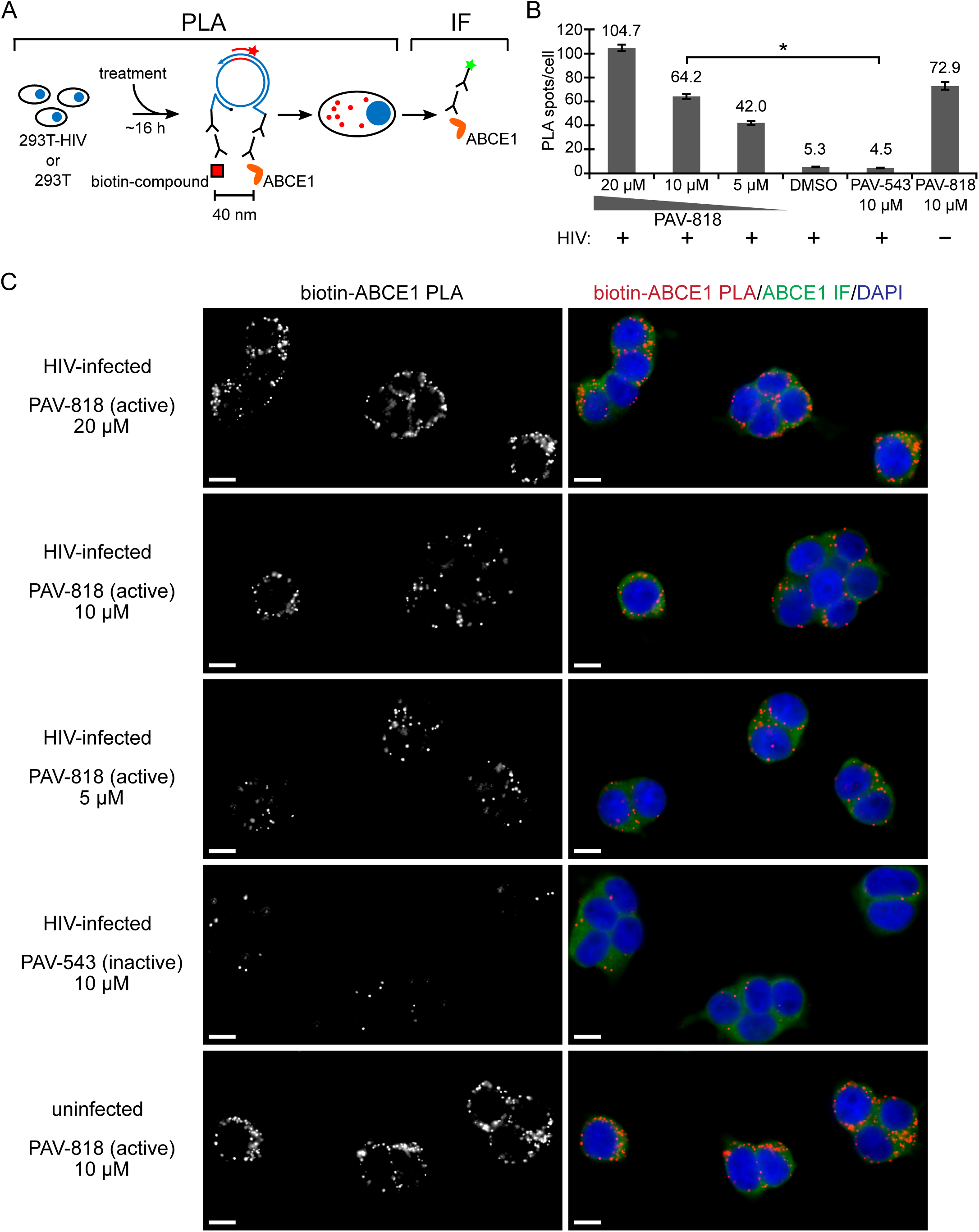
An antiviral PAV-206 analog colocalizes *in situ* with ABCE1, a host component of assembly intermediates. (A) Schematic of the PLA approach for detecting colocalization of compound with ABCE1. 293T cells chronically infected with HIV-1 (293T-HIV) or uninfected 293T cells were treated with indicated amounts of PAV-818 (the biotinylated active compound), PAV-543 (the biotinylated inactive compound), or DMSO for 16 h. PLA was performed by incubating with primary antibodies (mouse anti-biotin and rabbit anti-ABCE1) followed by PLA secondary antibodies (anti-mouse IgG coupled to (-) PLA oligo and anti-rabbit IgG coupled to (+) PLA oligo). Addition of other PLA reagents leads to connector oligos linking the (+) and (-) oligos only if the primary antibodies are colocalized; this in turn results in the PLA amplification reaction. Addition of an oligo that recognizes a sequence in the amplified regions and is coupled to a red fluorophore (red star) results in intense red spots at sites where biotinylated compound and ABCE1 are colocalized *in situ*. Following PLA, indirect immunofluorescence (IF) is performed by adding secondary antibody conjugated to a green fluorophore (green star) to detect any unoccupied ABCE1 antibody, thus marking intracellular ABCE1 with low-level green fluorescence (B) The graph shows the average number of biotin-ABCE1 PLA spots per cell for each condition, with + indicating HIV-1-infected cells and – indicating uninfected cells. Ten fields were analyzed for each group (containing a total of 104 - 155 cells per group), with error bars showing SEM. * indicates a significant difference in the number of biotin-ABCE1 PLA spots per cell when comparing treatment with PAV-818 vs. PAV-543, both at 10 μM (p < 0.001). (C) A representative field for each group quantified in (B) is shown, except for DMSO treatment. Fields on the left show biotin-ABCE1 PLA spots alone in grayscale. To the right are the same fields shown as a merge of three color-53 channels: biotin-ABCE1 PLA (red), ABCE1 IF (green), and DAPI-stained nuclei (blue). Scale bars, 10μm.

**FIG. 9.**
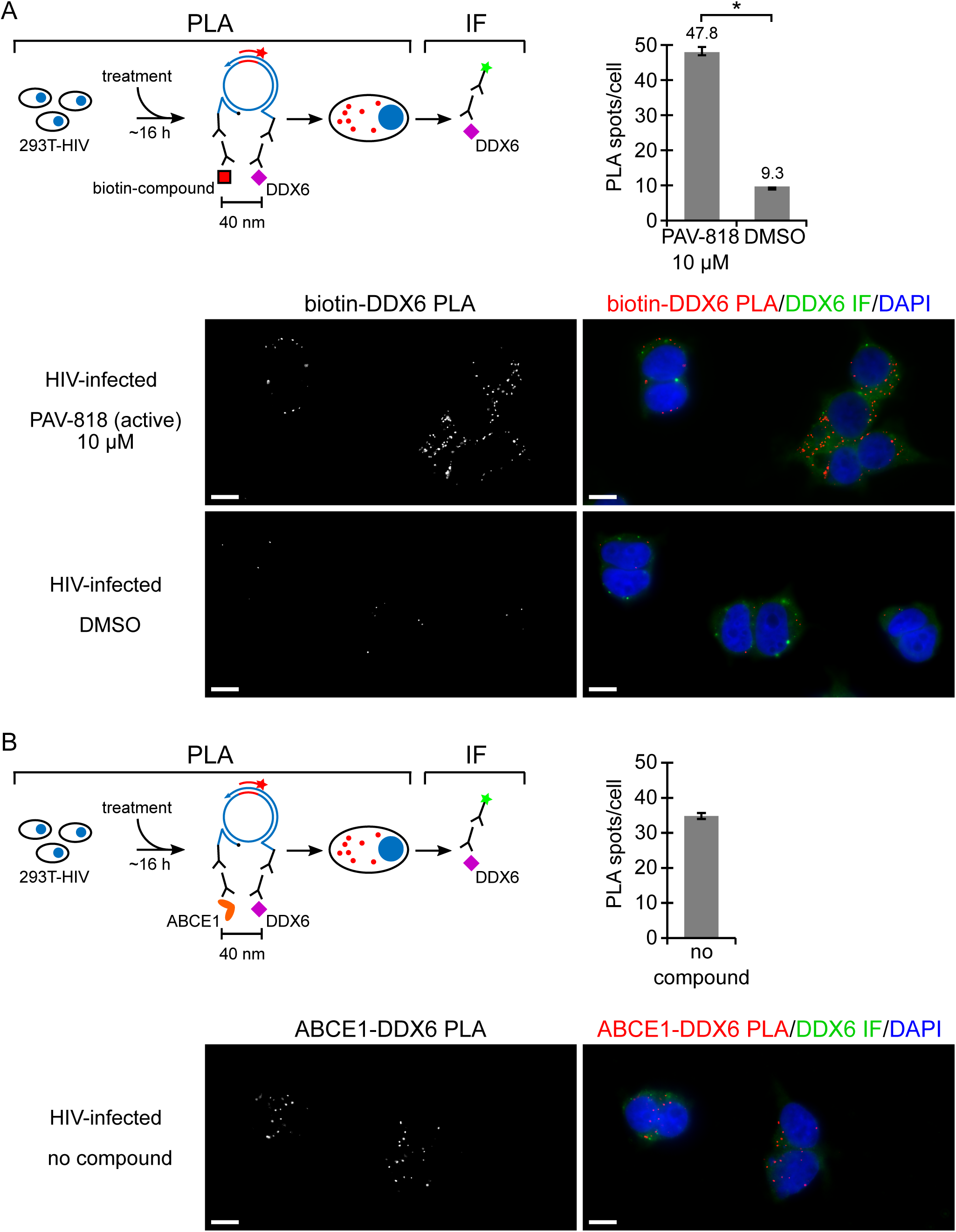
An antiviral PAV-206 analog colocalizes *in situ* with DDX6, an RNA granule protein in assembly intermediates that colocalizes with ABCE1. (A) A schematic of the PLA approach for detecting colocalization of compound with DDX6 is shown. 293T cells chronically infected with HIV-1 (293T-HIV) were treated with 10 μM PAV-818 (the biotinylated active compound) or DMSO for 16 h. PLA was performed by incubating with primary antibodies (mouse anti-biotin and rabbit anti-DDX6) followed by PLA secondary antibodies (anti-mouse IgG coupled to (-) PLA oligo and anti-rabbit IgG coupled to (+) PLA oligo). Addition of other PLA reagents leads to connector oligos linking the (+) and (-) oligos only if the primary antibodies are colocalized; this in turn results in the PLA amplification reaction. Addition of an oligo that recognizes a sequence in the amplified regions and is coupled to a red fluorophore (red star) results in intense red spots at sites where biotinylated compound and DDX6 are colocalized *in situ*. Following PLA, indirect immunofluorescence (IF) is performed by adding secondary antibody conjugated to a green fluorophore (green star) to detect any unoccupied DDX6 antibody, thus marking intracellular DDX6 with low-level green fluorescence. The graph shows the average number of biotin-DDX6 PLA spots per cell for each condition. Five fields were analyzed for each group (containing a total of 60 - 73 cells per group), with error bars showing SEM. * indicates a significant difference in the number of biotin-DDX6 PLA spots per cell when comparing treatment with PAV-818 vs. DMSO (p < 0.001). A representative field for each group quantified in the graph is shown, with images on the left displaying biotin-DDX6 PLA spots alone in grayscale and images on the right displaying a merge of three color-channels: biotin-DDX6 PLA (red), DDX6 IF (green), and DAPI-stained nuclei (blue). Scale bars, 10 μm. (B) A schematic of the PLA approach for detecting colocalization of ABCE1 with DDX6 is shown. 293T cells chronically infected with HIV-1 (293T-HIV) but not treated with any compounds were analyzed by PLA, as described for (A) above, except the primary antibodies used were rabbit anti-ABCE1 and mouse anti-DDX6, with red spots representing sites where ABCE1 and DDX6 are colocalized *in situ*. The graph shows the average number of ABCE1-DDX6 spots per cell. Ten fields were analyzed (containing a total of 121 cells), with error bars showing SEM. A representative field is shown, with the image on the left displaying ABCE1-DDX6 PLA spots alone in grayscale and the image on the right displaying a merge of three color-channels: ABCE1-DDX6 PLA (red), DDX6 IF (green), and DAPI-stained nuclei (blue).

## DISCUSSION

Here we report our discovery of a family of small molecules, PAV-206 and its analogs, that potently inhibit HIV-1 replication by acting on intracellular viral late events, a stage of the viral life cycle that is not specifically targeted by potent inhibitors currently in use or reported to be in development. Importantly, the exact target and mechanism of action of this compound remain to be determined; however, the data from virologic, resistance, and imaging studies presented here raise the intriguing possibility that this compound does not target a viral protein, but instead may target a host component of a multi-protein complex that plays an important role in assembly of the immature HIV-1 capsid (Fig. 10). While further studies will be needed to test this hypothesis, PAV-206 and its analogs should be of great interest even at this early stage given that all antiretroviral drugs in use except one (CCR5 antagonists) target viral proteins and that host-targeting antivirals offer a high genetic barrier to resistance that will be key as resistance to existing drugs becomes more prevalent (55, 56).

**FIG. 10.**
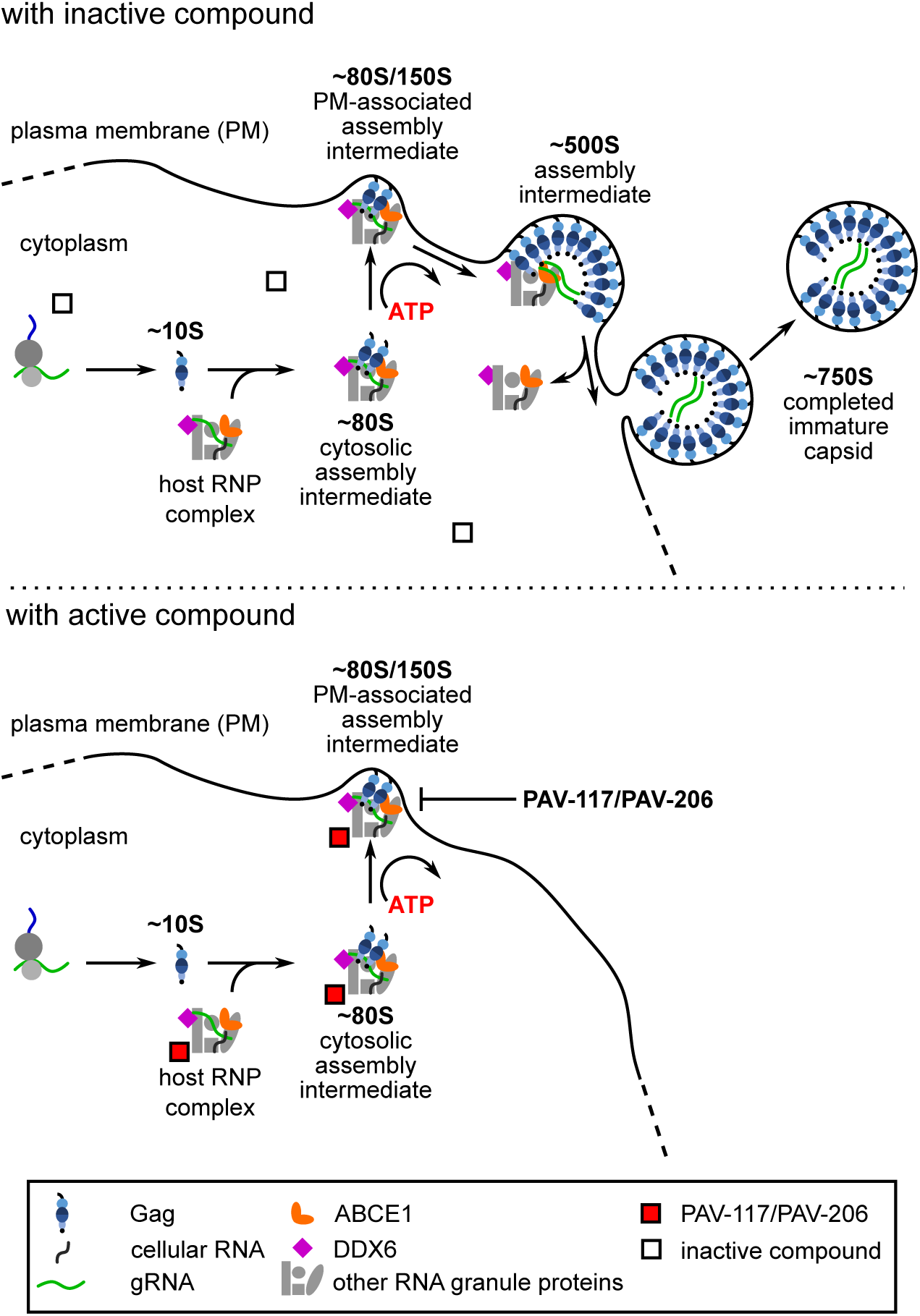
A model for the action of PAV-206, a possible first-in-class selective inhibitor of HIV-1 assembly. Upper diagram shows the intracellular pathway for HIV-1 immature capsid assembly, in the presence of an inactive compound, starting with synthesis of HIV-1 Gag and formation of the early ∼10S assembly intermediate. The ∼80S assembly intermediate is formed when HIV-1 Gag co-opts a host RNP complex related to an RNA granule to form the ∼80S cytosolic intermediate. This and subsequent intermediates contain Gag associated with ABCE1, DDX6, other host factors, and HIV-1 gRNA. After targeting of the cytosolic ∼80S assembly intermediate to the host plasma membrane, Gag multimerization continues leading to the formation of the ∼500S assembly intermediate. The host proteins are released upon formation of the ∼750S completed immature capsid, and budding results in release of the completed immature capsid. The lower diagram shows one model for PAV-117/PAV-206 activity based on the findings in this study. Our colocalization studies suggest that the compound is associated with both the HIV-1 assembly intermediates and the host RNA-granule-related complex from which they are derived, since we find the compound colocalized with a host component of this complex, ABCE1, in the presence and absence of Gag. Additionally, we observed a reduction of virus production and the amount of ∼500S assembly intermediate in PAV-117-treated infected cells. Thus, we hypothesize that PAV-117/PAV-206 inhibits virus production by acting during the HIV-1 immature capsid assembly pathway to inhibit progression past the ∼80S/150S assembly intermediate.

Most exciting in this regard was our observation that after selection with increasing concentrations of PAV-206 for 37 weeks in MT-2 T cells, sequence of replicating virus revealed no compound-specific mutations in *gag* or *pol* (Fig. 7), suggesting that PAV-206 acts by a mechanism that is independent of direct targeting of Gag. Additionally, high level resistance to PAV-206 was not observed during this time (Fig. 7). Such behavior is distinctly unusual, since all existing classes of direct-acting antiretroviral drugs result in rapid development of resistance mutations in their viral targets upon single drug treatment (41). The unusual nature of this finding was emphasized by our contrasting finding that 37-week selection with the protease inhibitor nelfinavir in parallel did result in three known resistance mutations in the HIV-1 protease as well as high-level resistance (allowing growth in drug concentrations 16X higher than the EC50), as would be expected.

We did observe two other categories of mutations during both PAV-206 and nelfinavir selection. The first are known tissue-culture adaptations (Fig. 7; Table S1 and S2). Most of these were in the cyclophilin A loop of Gag and are known to reduce the amount of cyclophilin A incorporated into virions during passage in cyclophilin A-rich immortalized CD4+ T cells, in the presence or absence of antiretroviral drugs (49, 50). One known tissue culture adaptation (54) was observed in *env* (Q550H) and arose in both selection groups. Mutations in a second category were also not compound-specific, and likely confer global replication advantage, as best demonstrated for A541V (51). The emergence of the earliest of these various compound-independent mutations, the cyclophilin A binding loop polymorphism V218M, as a subdominant mutation might explain the modest resistance observed in the PAV-206 selection experiment (allowing virus replication in a PAV-206 concentration 8X higher than the EC50). In support of this possibility, the appearance of modest resistance (to 8X EC50) in the nelfinavir selection control by passage 20 correlated with emergence of a cyclophilin A binding loop polymorphism and preceded the time when the drug-specific protease mutations became dominant. Further studies will be needed to test the hypothesis that these mutations allow low level replication in the presence of compound.

While additional studies will be required to identify the target of PAV-206, the surprising lack of compound-specific viral resistance mutations in *gag* or *pol* raises the possibility that this compound acts on a host protein. Two other observations are consistent with that hypothesis. First, this compound was identified using novel screens that recapitulate host-catalyzed pathways in which assembling capsid proteins form a series of host-protein containing assembly intermediates, culminating in formation of the completed immature capsid. Such an assembly pathway screen was previously shown to identify a host-targeting small molecule that potently inhibits RABV in cell culture (35). Second, our imaging studies show that PAV-206 colocalizes with Gag as well as ABCE1 and DDX6 (Figs. 6, 8, 9), two host enzymes known to be associated with Gag in assembly intermediates as shown by PLA and coimmunoprecipitation (24-27, 30, 32-34); here, we additionally showed that ABCE1 colocalizes with DDX6 by PLA (Fig. 9), consistent with previous co-immunoprecipitation results (30). Interestingly, our studies found that PAV-206 colocalizes with at least one of these host enzymes (ABCE1) in the absence of HIV-1 infection (Fig. 8), suggesting that the compound acts on the host RNA-granule-related RNP complex that is the precursor of assembly intermediates, even in the absence of Gag. Additional studies will be required to identify the direct target of this chemotype through crosslinking approaches that utilize the diazirine group in PAV-818, but based on data presented here, we hypothesize that the target may be a component of the host-RNP-complex-derived assembly intermediates.

PAV-206 and its analogs are also of great interest because, to our knowledge, they are the first potent small molecules shown to selectively inhibit late events in the HIV-1 life cycle. Unlike previously reported compounds that target CA with high potency (i.e. EC50 in the nM range or below, e.g. (12, 17, 57–59)) the chemotype reported here has no effects on viral early events and acts entirely on late events (Fig. 3). Moreover, PAV-117 dramatically reduces virus production (Figs. 3, 4), indicating that it acts on intracellular late events. PAV-117 treatment of infected cells did not reduce steady-state levels of two cellular proteins suggesting that PAV-117 does not globally inhibit protein synthesis as would be expected based on the screening strategy. PAV-117 treatment did lead to a modest reduction in steady-state Gag levels, although it is unlikely this modest reduction in steady-state Gag explains the reduction in virus production. Although we cannot rule out the possibility that the modest reductions in steady-state Gag levels are due to PAV-117 specifically inhibiting Gag synthesis, we favor the hypothesis that the reduced Gag levels are due to degradation of assembly intermediates that are associated with compound. Consistent with this hypothesis we do observe a reduction in the steady-state levels of the final (∼500S) assembly intermediate (Fig. 4). Nevertheless, additional studies will be needed to determine the exact mechanism of action of PAV-206 and its analogs.

Finally, it should be noted that discovery of these compounds constitutes a compelling validation of the importance of HIV-1 RNA-granule-related assembly intermediates in virus production. Surprisingly, the existence of assembly intermediates containing RNA granule proteins continues to be questioned, even though numerous groups in addition to ours have implicated RNA granule proteins as being important for assembly of diverse retroviruses and retroelements. For example, knockout studies by the Sandmeyer group demonstrated that RNA granule proteins are required for Ty3 retrotransposition in yeast and likely act during assembly (60–62). Additionally the Linial group found that the RNA granule protein DDX6 is required for foamy virus packaging (63), and the Mouland group has shown that the RNA granule protein Stau1 plays an important role in HIV-1 Gag assembly and packaging (64–67). Finally, the Kewalramani and Pathak groups have demonstrated that the RNA granule protein MOV10 influences immature HIV-1 assembly (68, 69). Data presented here shows that the study of host-derived RNA-granule-derived HIV-1 assembly intermediates has led to discovery of the first compounds that specifically inhibit intracellular HIV-1 late events. Thus, these data add strong support to the view that such assembly intermediates not only exist but are highly relevant to our understanding of HIV-1 assembly as well as development of antiretroviral compounds for treatment of multidrug resistant HIV-1 in the future.

## METHODS

### Plasmids and proviral constructs

In this study several variants of the HIV-1 NL4-3 proviral clone were utilized. For infection of PBMC, we utilized the NL4-3 Infectious Molecular Clone (pNL4-3) from Dr. Malcolm Martin obtained through the NIH AIDS Reagent Program, Division of AIDS, NIAID, NIH (Catalog Number 114; (70)). For infection of MT-2, the pNL4-3 plasmid was modified to express *Renilla* luciferase in place of *nef* to make the HIV-1 pNL4-3 RLuc plasmid. For single round MT-2 cell infections used to assay only HIV-1 early events, we introduced an *env* deletion (KpnI-BglII) into the RLuc plasmid to make pNL4-3 RLuc Δ*env*. We also used several variants of the HIV-1 LAI proviral clone. Chronically infected H9 and 293T cells were generated using HIV-1 pLAI *vp*r-*nef*-*puro*+, a proviral plasmid that contains a puromycin gene in place of *nef* for selection and two stop codons introduced into the NcoI site within *vpr* to inactivate Vpr (71). For single round MT-2 cell infections used to evaluate viral late events, we used pLAI *pro*-Δ*env* which we have described previously (30). The plasmid pcDNA NL4-3 Env used for pseudotyping was generated by cloning a Geneblock DNA fragment of codon-optimized NL4-3 *env* (Integrated DNA Technologies ((IDT), Coralville, IA) into a pcDNA vector by Gibson assembly (72). For infection of SupT1.CCR5 cells, we utilized a molecular clone of a subtype A primary isolate, Q23-17(73), that was modified to express a subtype A BG505 *env* (74, 75) and was obtained from Julie Overbaugh. Infection of G355-5 cells with FIV utilized the FIV-34TF10 ORFA+ proviral clone, described previously (31). The psPAX2 Gag-Pol helper plasmid was obtained from Didier Trono via Addgene (plasmid # 12260; http://n2t.net/addgene:12260). To evaluate compound activity against FLUV, plasmids required for production of FLUV A/WSN/33 (76) were obtained from Andrew Pekosz.

#### Cells and transfection

The feline astrocyte cell line G355-5 was obtained from ATCC (Manassas, VA; CRL-2033) and maintained in McCoy 5A (modified) medium supplemented with 10% fetal bovine serum (complete McCoy 5A). G355-5 cells were transfected with 1 to 6 µg DNA in 6 cm dishes using X-treameGENE 9 (Roche, Basel, Switzerland). The 293T/17 cell line was obtained from ATCC (CRL-11268) and maintained in DMEM media supplemented with 10% fetal bovine serum and 1% penicillin-streptomycin (complete DMEM). 293T cells were transfected with 1 mg/mL polyethylenimine (PEI; Polysciences, Warrington, PA) at a ratio of PEI-to-DNA of 3:1. The MT-2 cell line was obtained from Dr. Douglas Richman through the NIH AIDS Reagent Program, Division of AIDS, NIAID, NIH (Catalog Number 237; (77, 78)). SupT1.CCR5 cells, a SupT1 cell line stably expressing CCR5 HIV-1 coreceptor (79),, were obtained from Julie Overbaugh. MT-2 and SupT1.CCR5 cells were maintained in RPMI supplemented with 10% fetal bovine serum and 1% penicillin-streptomycin (complete RPMI). TZM-bl cells were obtained from Dr. John C. Kappes, and Dr. Xiaoyun Wu through the NIH AIDS Reagent Program, Division of AIDS, NIAID, NIH (Catalog number 8129; (80–84)). The MDCK.2 cell line was obtained from ATCC (CRL-2986) and were maintained in MEM media supplemented with 10% fetal bovine serum, 1% penicillin-streptomycin, 1% non-essential amino acids solution, and 1% sodium pyruvate (complete MEM).

#### HIV-1 chronically infected cell lines

The 293T-HIV and H9-HIV lines stably express the provirus LAI *vpr-nef-puro+*. These lines were generated by infecting either 293T/17 or H9 cells with NL4-3 *vpr-nef-puro+* at an MOI of 1, followed by selection with and maintenance in complete media supplemented with 0.5 μg/mL puromycin.

#### Antibodies for western blotting and proximity ligation assay

To detect HIV-1 Gag p24 in cell lysates and virus pellets by WB, we used αGag p24 mouse IgG_1_ monoclonal antibody isolated from the hybridoma 183-H12-5C obtained from Bruce Chesebro through the AIDS Reagent Program Division of AIDS, NIAID, NIH (catalog number 1513; (85)). Commercial antibodies were used to detect GAPDH (Abcam, Cambridge, MA; ab9485) and actin (Millipore-Sigma, St Louis, MO; A2066) in cell lysates by WB. Isotype specific secondary antibodies utilized for WB were obtained from Bethyl Laboratories (Montgomery, TX). For proximity ligation assay (PLA) we utilized the αGag antibody described for WB above, αbiotin rabbit IgG (Bethyl Laboratories, A150-109A), affinity purified αABCE1 rabbit IgG which we have previously described (34), αbiotin mouse IgG (Jackson Immunoresearch Labs, West Grove, PA,; 200-002-211), αDDX6 mouse IgG (Millipore-Sigma; WH0001656M1), and αDDX6 rabbit IgG (Bethyl Laboratories; A300-461A). Rabbit IgG (Millipore-Sigma; I5006) and mouse IgG (Millipore-Sigma; I5381) was used for PLA nonimmune controls.

#### Virus infectivity measurement

Infectivity was measured by titering stocks on TZM-bl cells. Cells seeded 7500 cells/well into 96-well polystyrene tissue culture plates were infected 24h later with dilutions of culture supernatant or virus stock containing 20 µg/ml DEAE dextran in a 50 µL volume for 4 h at 37°C. After infection, 150 µL of complete DMEM was added and plates were incubated at 37°C for 48 h. Cells were fixed with 1% formaldehyde/0.2% glutaraldehyde/PBS for 5 min and washed with PBS. To determine infectious units, cells were incubated with 50 µL of X-Gal stain (4 mM potassium ferrocyanide, 4 mM potassium ferricyanide, 2 mM MgCl_2_, 0.4 mg/mL 5-Bromo-4-Chloro-3-Indolyl β-D-Galactopyranoside prepared in PBS) for 50 min at 37°C. Following staining cells were washed several times with PBS and blue cells (only wells with < 100 cells) were counted to determine infectious units (IU) per mL. To determine relative infectivity, cells were seeded, infected, fixed, and washed as above, but instead of X-Gal stain, 100 µL of MUG stain (200 µg/mL 4-methylumbelliferone β-D-galactopyranoside prepared in DMEM) was added and incubated at 37°C for 25 min. After incubation, the reaction was terminated with 100 µL of 1M Na_2_CO_3_ and well fluorescence determined using 360 nm and 449 nm for excitation and emission settings, respectively (Synergy H1; BioTek Instruments, Inc., Winooski, VT, USA). The relative fluorescence was used to compare infectivity between samples.

#### PBMC activation

Thawed, unstimulated PBMC cells (BenTech, Seattle, WA) were pelleted by centrifugation 228 x g for 8 min in an Allegra centrifuge (Beckman, Brea, CA). Pellets were resuspended in activation medium containing RPMI supplemented with 21 ng/mL human IL-2 (Peprotech, Rocky Hill, NJ; #200-02), 1.5 μg/mL PHA (Remel, San Diego, CA; R30852801), 10% fetal bovine serum, and 1% penicillin-streptomycin, at a density of 2×10^6^ cells/mL. After a three-day incubation, cells were collected and resuspended in RPMI with 20% FBS and 10% DMSO, and frozen as 50×10^6^ cells per vial.

#### HIV-1 virus production

To produce HIV-1 viral stocks 293T/17 cells were seeded at 10×10^6^ cells in a T175 flask, transfected for 24 h, washed, and incubated in complete DMEM. Media from 48 and 72 h post transfection collections was pooled, filtered with a 0.45 µm PES membrane, and 0.5 mL aliquots were stored at −80°C. To produce viral stocks of NL4-3, Q23.BG505 and LAI *vpr*-*nef*-*puro*+, 36 µg of proviral plasmids were used per T175 flask. To produce viral stocks of NL4-3 RLuc Δ*env* pseudotyped with NL4-3 Env, 36 µg of the proviral plasmid, and 12 µg of pcDNA NL4-3 Env were co-transfected. To produce viral stocks of LAI pro-Δ*env* pseudotyped with NL4-3 Env, 27 µg of proviral plasmid, 15 µg of pcDNA NL4-3 Env, and 18 µg of psPAX2 (Gag-Pol helper plasmid) were co-transfected.

#### FLUV virus production

FLUV A/WSN/33 viral stocks were prepared in MDCK.2 cells as described previously (86).

#### FIV virus production and RT assay

G355-5 cells were transfected in 6-well plates and each well was transfected with 2 µg per well of FIV 34TF10 ORFA+ proviral plasmid. At 24 h post transfection, cells were washed and media replaced with 4 ml of complete McCoy 5A. At 144 h post transfection, media was filtered with a 0.45 µm syringe filter, aliquoted, and stored at −80C. Reverse transcriptase activity in media from FIV or mock-transfected cells was estimated using SG-PERT (87). A set of recombinant RT standards (EMD Millipore, Molsheim, France; catalog# 382129) ranging from 5.12e3 nU/mL to 5.12e9 nU/mL was used to estimate the nU/mL of reverse transcriptase activity in the FIV stock.

### Inhibition of HIV-1 replication

*MT-2 spreading infection assay*. Inhibition of HIV-1 viral replication was assayed in a spreading infection using MT-2 cells and NL4-3 RLuc reporter virus. For dose-response curves, compounds were initially diluted with DMSO to 100-times the starting concentration in a 96-well plate and subjected to a series of 3-fold dilutions in DMSO for a total of eight or nine dilutions. If a single compound concentration was tested, the compound was diluted to 100-times the desired concentration in DMSO. Compounds were then diluted 50-fold with infection media prepared by diluting NL4-3 RLuc virus stock to 400 IU/100 μL with complete RPMI. Then 100 μL of 50-fold diluted compound was transferred to 20,000 MT-2 cells that were pre-seeded in 96-well plates in 100 uL complete RPMI for a final volume of 200 u, followed by incubation at 37°C for 96 h. The final MOI in infected plates was 0.02 and the final DMSO concentration in all wells was 1%. All assays were run with three replicates. For each replicate, one well received DMSO only and one well received media only for normalization and background correction. To assay inhibition of HIV-1 replication, 100 μL of media was removed and discarded and 10 μL of 15 μM EnduRen luciferase substrate was added to each well, followed by incubation for 1.5 h at 37°C. Plates were read on a luminescence plate reader (Synergy H1; BioTek Instruments, Inc.).

*SupT1.CCR5 spreading infection assay*. The SupT1.CCR5 assay was as described for the MT-2 spreading infection assay, with the following exceptions: 12.5 uL of Q23.BG505 virus stock was used per well, and inhibition of HIV-1 replication was determined by measuring the relative infectivity produced form infected cells using the MUG assay (see virus infectivity measurements above).

*PBMC spreading infection assay*. The PBMC assay was as described for the MT-2 spreading infection assay with the following exceptions. Frozen activated PBMC cells were thawed and resuspended in complete PBMC media containing RPMI supplemented with 21 ng/mL human IL-2 (Peprotech; #200-02), 10% fetal bovine serum and 1% penicillin-streptomycin, at 0.5×10^6^ cells/mL for 3 days in 37°C tissue culture incubator. Cells were collected and adjusted to 2×10^6^ cells/mL in complete PBMC media and bulk infected for 2 h with NL4-3 virus stock at a MOI of 0.008 at 37°C. Compound dilutions were prepared in complete PBMC media as above, but without virus, and 100 uL of diluted compound was transferred to 96-well U-bottom plates. Bulk infected cells were pelleted and resuspended in complete PBMC media at 1×10^6^ cells/mL and 100 μL of cell suspension was transferred to each well at 100,000 cells per well with diluted compound. After 96 h, media was collected for relative infectivity measurements as with the SupT1.CCR5 spreading infection assay.

#### FIV inhibition assay

Inhibition of FIV spreading infection was assayed using G355-5 cells, seeded on the prior day at 8000 cells/well in a 96-well plate. Preparation and addition of compounds were as in the MT-2 spreading infection assay with the following differences: for infection, 1954 nU RT activity of the FIV-34TF10 ORFA+ stock was used per well, infection assays proceeded for 144 h, and infectious virus production was measured using the SG-PERT assay described above.

#### Virus pelleting

For Fig. 3B, culture supernatants were collected at 48 h and virus was pelleted by adding 75 μL of 17% PEG 6000/0.8M NaCl/PBS to 75 μL media from the assay in a 96-well V-bottom polypropylene plate. PEG and media were incubated on ice for 2 h or overnight at 4°C. Virus was pelleted at 1847 x g for 30 min at 4°C in the plate in an Allegra centrifuge (Beckman) and the pellet was resuspended in 50 uL of loading buffer with DTT for analysis by WB with *α*Gag. For Fig. 4, culture supernatants were collected 48 h post infection and filtered through a 0.45 μm PES syringe filter to remove any remaining cells. Virus-like particles were centrifuged through a 30% w/v sucrose/PBS cushion at 130,000 x g, 30 min, 4°C in an SW55Ti rotor (Beckman), and the virus pellet harvested for analysis by WB with *α*Gag.

#### MT-2 acute infection assay

MT-2 cells were diluted to 2×10^6^ cells/mL with complete RPMI and bulk infected at a MOI of 1 for 2 h at 37°C. Compound was diluted to 100-times the final required concentration with DMSO. The compound dilutions and DMSO only controls were then further diluted 50-fold with complete RPMI, and 1 mL of the dilutions was added per well in a 12-well plate. Bulk infected cells were then washed and adjusted to 0.5×10^6^ cells/mL, and 1 mL of infected cells was added to each well containing diluted compound or DMSO. At 48h, media was collected for virus pelleting as described above and cells were washed and lysed (see cell lysis below).

#### Cell lysis

MT-2 cells were collected and washed three times by pelleting at 200 x g for 5 min at 4°C in an Allegra centrifuge (Beckman) and resuspending cells in ice-cold PBS. Cell pellets were lysed with 250 µL lysis buffer (20 mM HEPES pH 7.9, 14 mM KAc, 1 mM MgCl_2_, 0.3% NP40) followed by shearing 20-times with a 20-gauge needle. Cell lysate was then clarified 200 x g for 10 min at 4°C in an Allegra centrifuge (Beckman). Supernatant was transferred to a fresh tube and further clarified by centrifugation at 18,000 x g for 10 min at 4°C in a microfuge (Beckman), and supernatant transferred to a fresh tube.

#### Velocity sedimentation

To prepare gradients, 10%, 15%, 40%, 50%, 60%, 70%, and 80% w/v sucrose solutions were prepared with water and then each was diluted 1/10 with a 10x stock of the lysis buffer used to harvest cells (see cell lysis). For each gradient, ∼100 μL of cell lysate (∼100 μg of total protein) was layered on a 5-ml step gradient containing equivalent layers of each diluted sucrose solution prepared and subjected to velocity sedimentation in a Beckman MLS-50 rotor 217,000 x g for 45 min at 4°C. Gradients were fractionated from top to bottom, and pelleted material was harvested for WB. Aliquots of fractions and pellet were analyzed by SDS-PAGE, followed by WB with αGag. Gag in pellet, which likely represents denatured Gag, was not included in the quantification of the velocity sedimentation gradients and represented less than 10% of total Gag signal in the gradient. The method for estimating the migration of particles with different S values in gradients has been described previously (28).

#### Inhibition of HIV-1 early events

For Fig. 3A, the assay described under MT-2 spreading infection assay was used with the following exceptions: MT-2 cells were infected with 0.5 μL/well of HIV-1 NL4-3 RLuc Δ*env* virus pseudotyped with NL4-3 Env and luciferase was measured at 48 h instead of 96 h.

#### Inhibition of HIV-1 late events

For these experiments, we used H9-HIV, a chronically infected H9 T cell line. Compounds were diluted as described under MT-2 spreading infection above. To measure inhibition of infectious virus production (Fig. 3B, black graph), media were collected and relative infectivity of virus in the media was measured using the MUG assay (see virus infectivity measurements above). To measure inhibition of virus production (Fig. 3B, blue graph and blot), PEG pelleted virus (harvested as described in virus pelleting above) was analyzed by WB with *α*Gag.

#### FLUV antiviral assay

Inhibition of FLUV replication was assayed using infection of MDCK.2 cells. A day before the assay, MDCK.2 cells were seeded at 30,000 cells/well in complete MEM in a 96-well plate. The next day, seeded cells are washed with PBS and incubated for 1 h with 100 μL of FLUV A/WSN/33 stock diluted in VGM media containing 42 mM HEPES, 0.125% BSA, and 1 μg/mL TPCK-trypsin (final MOI of 0.001). Cells were then washed with PBS and incubated in 90 uL of complete MEM. Single compound dilutions were prepared in complete MEM at 10x the desired final concentration, and 10 μL of DMSO or 10x compound was added to infected cells for final volume of 100 μL. At 24 h post infection, media was aspirated, cells were washed with PBS, and 100 μL of complete MEM was added to each well. After a 2 h incubation, media were collected for TCID50 determination.

#### FLUV TCID50 assay

FLUV TCID50 was assessed on MDCK.2 cells. MDCK.2 cells were seeded at 30,000 cells/well in complete MEM in a 96-well plate and the next day the complete MEM was exchanged for 100 μL VGM media (as in the FLUV antiviral assay). Ten-fold serial dilutions of virus-containing supernatant were prepared and seven replicates of 11.1 μL of neat and diluted stocks were added to the MDCK cells (final dilutions in the TCID50 plate ranged from 10^-1^ – 10^-6^). Plates were incubated at 37°C for 3 days, and the number of infected wells in the seven replicates for each dilution was determined by visual inspection. These data were used to calculate TCID50/mL using the Reed and Muench method (88).

#### Cytotoxicity Assays

For each viral inhibition assay, a cytotoxicity assay was performed in parallel on uninfected cells by adding 1/10^th^ volume of AlamarBlue (Thermo Fisher, Waltham, MA; DAL1199) to the plates and mixing well. AlamarBlue-treated plates were incubated for 2-3 h at 37°C (timing was optimized for linear signal), mixed well, and read using a fluorescence plate reader with 560 nm and 590 nm as excitation and emission settings, respectively (Synergy H1; BioTek Instruments, Inc.).

#### Resistance Assay

Resistance selections were carried out in 12-well tissue culture plates. First, compound dilutions were prepared by adding 1.25 mL of complete RPMI to each well followed by 25 μL of compound that was diluted to 100x the desired final concentration in DMSO. For the first passage, NL4-3 virus stock was then added to achieve a MOI of 0.01. For subsequent passages, 150 μL of the previous passage was added instead. Finally, 0.2×10^6^ MT-2 cells were added in 1.25 mL of complete RPMI. The cultures were incubated at 37°C until they reach maximum viral CPE, as determined by visual inspection for the formation of syncytia (3-7 days). Selection was begun at 1x[EC50] and then increased 2-fold whenever maximum CPE was achieved in a time frame similar to that achieved by virus passaged in DMSO. At the end of each selection cells were pelleted from the media by centrifugation 200 x g for 5 min in an Allegra centrifuge (Beckman) and supernatant was transferred for storage at −80C. PAV-206 and nelfinavir selections were carried out in parallel for 37 weeks.

#### Sequencing virus from resistance selection

Passaged media was first titered on TZM-bl cells and infectious units/ mL determined by X-Gal staining (see Virus infectivity measurement above. Passaged virus was then used to infect MT-2 cells at a MOI of 1 in complete RPMI supplemented with 100 ug/mL DEAE dextran for 3 h at 37°C. Cells were pelleted at 200 x g 5 min in Allegra centrifuge (Beckman) and then resuspended in complete RPMI. At 15 hours post infection, cells were washed three times with ice-cold PBS and then viral cDNA from the cell pellet was harvested using a Spin Miniprep kit (Qiagen, Venlo, Netherlands). Viral cDNA was amplified by PCR using the Herculase II polymerase (Agilent, Santa Clara, CA) and amplified viral cDNA was gel purified using a Spin Gel Purification kit (Qiagen). Two sets of primers were used to amplify viral cDNA in two overlapping fragments. The first half of the genome from 132 base pairs before the start of *gag* to midway through *vpr* was amplified with the forward primer 5’-AAGCGAAAGTAAAGCCAGAGG-3’ and reverse primer 5’-AACGCCTATTCTGCTATGTCG-3’. The second half of the genome from *vpr* to 252 base pairs into *nef* was amplified with the forward primer 5’-CAGAGGACAGATGGAACAAGC-3’ and the reverse primer 5’AGCTGCCTTGTAAGTCATTGG-3’. The LTRs were not sequenced. Gel purified PCR products were then subjected to Sanger sequencing. Gel chromatograms were visually inspected to identify a second chromatogram peak that was equal or larger in height than the peak corresponding to the reference sequence (defined as a ≥ 50% mutation; see Table S1 and S2) or a single chromatogram peak that differs from the reference sequence (defined as a dominant mutation; see Table S1 and S2).

#### Proximity Ligation Assay (PLA)

For each treatment, one-half of a Grace Biolabs Culture Well silicone chamber (Millipore-Sigma; GBL103380) was attached to a cover glass (Carl Zeiss, Oberkochen, Germany; #474030-9000-000) to create four chambers on the cover glass. Each chambered cover glass was placed into a well of a 6-well plate and chambers were coated with poly-L-lysine and allowed to dry overnight. The next day ∼4×10^3^ 293T/17 or 293T-HIV cells were seeded into each chamber and incubated at 37°C for 5-6 h. After cells adhered, they were treated with compound or DMSO. At 16 h post treatment, cells were fixed for 15 min in 4% paraformaldehyde in PBS pH 7.4, permeabilized in 0.5% w/v Saponin in PBS, pH 7.4 for 10 min, and blocked in Duolink blocking solution (Millipore-Sigma) at 37°C for 30 min. Cells were incubated in primary antibodies using the following concentrations: for the Gag-Biotin PLA, mouse αGag p24 was used at 0.2 ug/mL and rabbit αbiotin was used at 4 ug/mL; for the ABCE1-Biotin PLA, rabbit αABCE1 was used at 0.3 ug/mL and mouse αbiotin was used at 0.7 ug/mL; for the DDX6-Biotin PLA, rabbit αDDX6 was used at 1 ug/mL and mouse αbiotin was used at 0.7 ug/mL; for the ABCE1-DDX6 PLA, rabbit αABCE1 was used at 0.3 ug/mL and mouse αDDX6 was used at 0.3 ug/mL. For the nonimmune (NI) control PLA experiments, NI control antibodies from the relevant species were used at the same concentration as the primary antibody they replaced. Primary incubation was followed by incubation with Duolink reagents (Millipore-Sigma): oligo-linked secondary antibody, ligation mix, and red amplification/detection mix, with washes in between, as per the Duolink protocol. For concurrent immunofluorescence (IF) following the final 1x PLA Buffer B washes, cells were incubated for 30 min at RT with 1:1000 Alexafluor 488 anti-mouse secondary antibody. Cover glasses were mounted using Duolink In Situ Mounting Media with DAPI (Millipore-Sigma), sealed to the glass slides with clear nail polish, allowed to dry for 24 h at RT, and stored at −20°C. Imaging was performed with a Zeiss Axio Observer.Z1/7 deconvolution microscope using Zeiss Plan-Apochromat 63X/ aperture 1.4 objective with oil immersion, with Zen 2.6 (blue edition) software. For quantification, five to ten fields, each containing at least three IF-positive cells, were chosen at random and imaged using identical exposure times for each channel (100 milliseconds for the PLA channel in red; 0.5-1 second for the IF channel in green, depending on IF primary antibody; and 20 milliseconds for the DAPI channel in blue). Images were captured as a Z-stack of 80-90 0.24-µm slices that included the entire cell body and 5 µm above and below the cell (∼20 µm total). To determine the number of PLA spots per cell, image stacks were processed as follows in FIJI (89). First, an in-focus slice from the IF channel was selected using the Find Focused Slices plugin within ImageJ (Tseng, Q (2020), available from https://sites.google.com/site/qingzongtseng/find-focus) and thresholding was used to generate regions of interest (ROI) that contained only cells. The ROIs identified in the IF channel were applied to a maximum intensity Z-projection of the PLA channel and PLA spots within the ROI were counted using the find maxima function in Fiji, with noise threshold optimized to identify only PLA spots. Nuclei from each field were counted manually to determine the number of cells. Total PLA spots counted were divided by total nuclei to obtain the average number of PLA spots per cell, and these results were plotted with error bars representing SEM, with *n* being the total number of cells counted. The number of fields and cells analyzed for each PLA were: Gag-biotin PLA (Fig. 6) 20 fields, containing a total of 186-316 cells; ABCE1-biotin PLA (Fig. 8) 10 fields, containing a total of 104 - 155 cells; DDX6-biotin PLA (Fig. 9) 5 fields, containing a total of 60 - 73 cells; DDX6-ABCE1 PLA (Fig. 9) 10 fields, containing a total of 121 cells; Gag-biotin PLA (Fig. S2) 20 fields, containing a total of 213 - 304 cells; rabbit biotin-mouse NI PLA (Fig. S2) 10 fields, containing a total of 107 - 118 cells; rabbit NI-mouse Gag PLA (Fig. S2) 10 fields, containing a total of 131 cells; ABCE1-biotin PLA (Fig. S3) 5 fields, containing a total of 53-66 cells; mouse biotin-rabbit NI PLA (Fig. S3) 5 fields, containing a total of 51 - 59 cells; mouse NI- rabbit ABCE1 (Fig. S3) 5 fields, containing a total of 44 - 77 cells. For presentation of images in figures, a central slice of the image stack was chosen, and a region of the field of 1875 pixels x 938 pixels was selected and adjusted as follows. For all channels, background was removed by setting the intensity minimum above the background signal. For the PLA channel, the gain on PLA spots was increased for visualization by setting the channel intensity maximum to approximately one-third the intensity of the most intense pixel in the selected field. For the IF channel, IF signal was reduced to improve PLA spot visualization by setting the channel intensity maximum to approximately twice the intensity of the most intense pixel in the selected field. A gray scale image of the PLA channel only and an RGB image of the merged channels (PLA, red; IF, green; and DAPI, blue) were generated and imported into Inkscape, an open source vector graphics editor, to create the final figure layout without further adjustment.

#### Statistical Analysis

EC50 and CC50 values were determined in Excel using four-parameter logistic regression analysis of dose-response data generated from the viral inhibition or cytotoxicity assays, respectively. Solid-lines on dose-response plots represent the fitted logistic regression. Average EC50 and CC50 values were calculated using the geometric mean and average values were reported with the geometric standard deviation (GSD). The statistical difference between pairs of treatment groups in the PLA experiments was analyzed in Excel using a two-tailed unpaired Students *t* test assuming unequal variance.

#### Synthesis of compounds

See Supplementary Methods.

## ACKNOWLEDGMENTS

The following reagents were obtained through the NIH AIDS Reagent Program, Division of AIDS, NIAID, NIH: αGag, p24 mouse IgG_1_ monoclonal antibody isolated from the hybridoma 183-H12-5C obtained from Bruce Chesebro (catalog number 1513); HIV-1 NL4-3 Infectious Molecular Clone (pNL4-3) from Malcolm Martin (Catalog Number 114); MT-2 cell line from Douglas Richman (Catalog Number 237); and TZM-bl cells from John C. Kappes and. Xiaoyun Wu (Catalog number 8129). HIV-1 Q23.BG505 virus and SupT1.CCR5 cells were obtained from Dr. Julie Overbaugh. The psPAX2 Gag-Pol helper plasmid was obtained from Didier Trono via Addgene (plasmid # 12260). Plasmids for production of FLUV A/WSN/33 were obtained from Andrew Pekosz.

We acknowledge Michael Emerman for advice and input.

Jaisri Lingappa is a cofounder of Prosetta Biosciences and Jonathan Reed is a consultant for Prosetta Biosciences. Vishwanath Lingappa is a cofounder, CEO, and CTO of Prosetta Biosciences.

These studies were funded by NIH R01AI150457, NIH R56 AI145498, and Prosetta Biosciences. Neither of the funders (NIH or Prosetta Biosciences) had any role in study design, data collection and interpretation, or the decision to submit the work for publication.

## SUPPLEMENTARY FIGURE LEGENDS

**FIG. S1.**
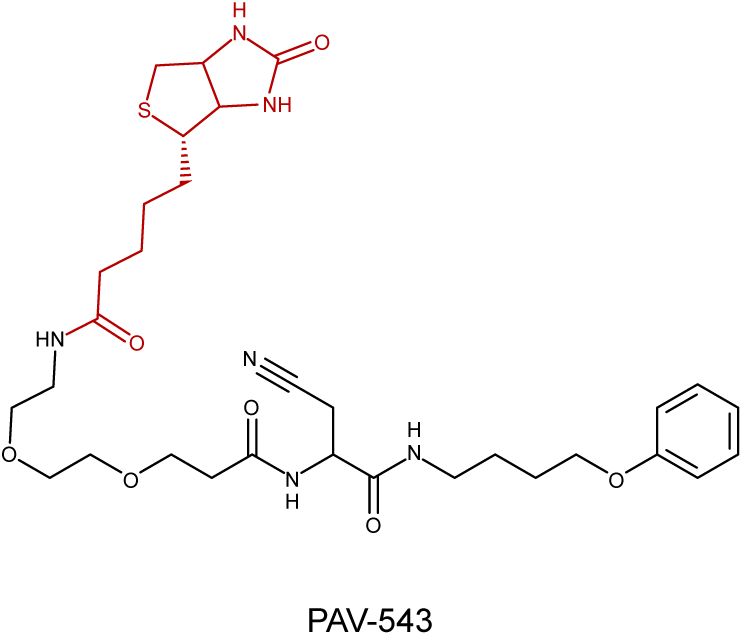
Structure of PAV-543, a biotinylated compound that does not have antiretroviral activity. The biotin moiety is shown in red. The EC50 of PAV-543 is > 100 μM in the HIV-1 MT-2 cell spread assay shown in Fig 2C as is the CC50 in MT-2 cells (Reed and Lingappa, unpublished observations), indicating no significant antiretroviral activity.

**FIG. S2.**
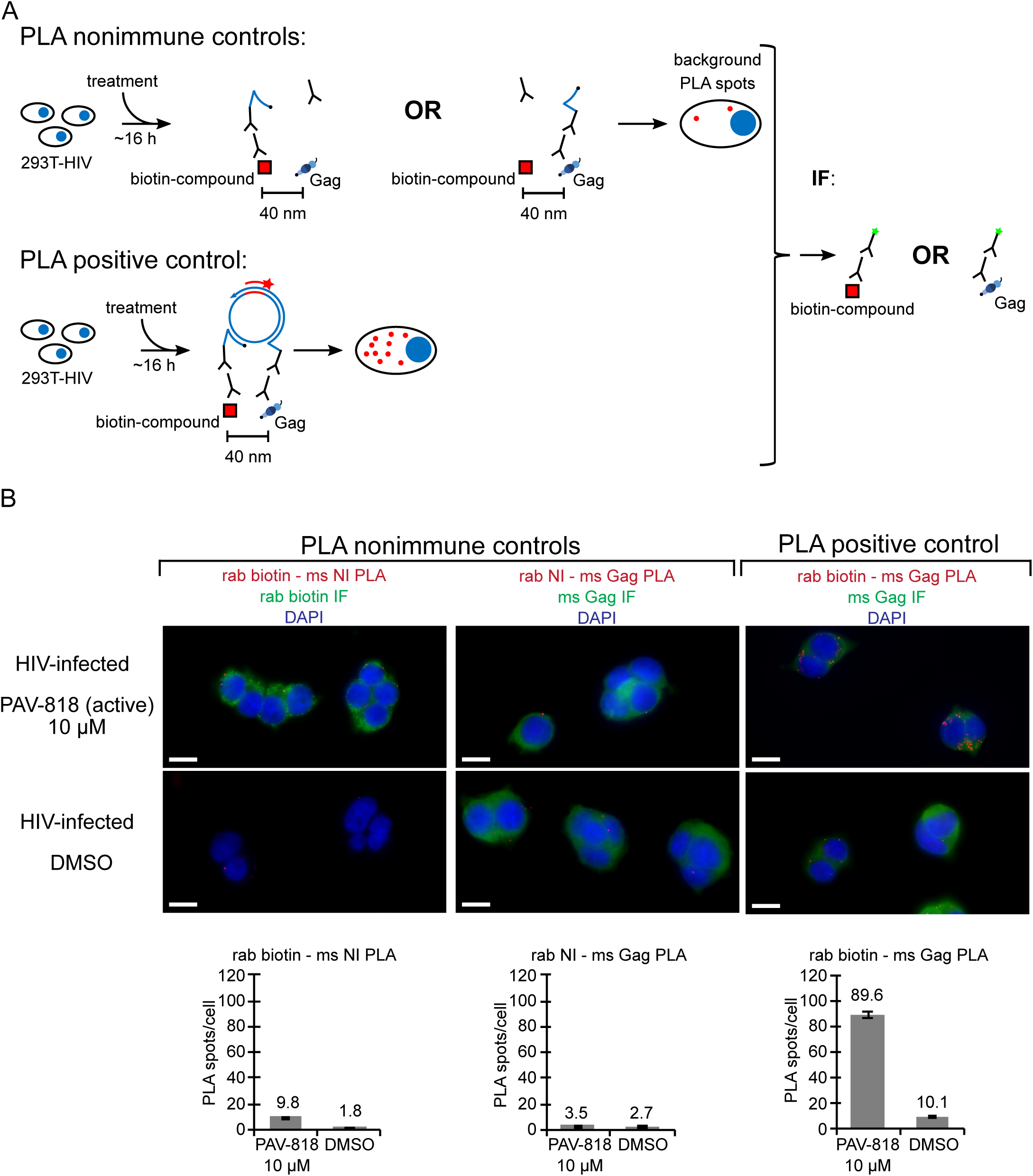
Nonimmune controls for the biotin-Gag proximity ligation assay. (A) Schematic of the PLA approach for testing specificity of the antibodies used to determine colocalization of compound with Gag. 293T cells chronically infected with HIV-1 (293T-HIV) were treated with 10 μM PAV-818 (the biotinylated active compound) or DMSO for 16 h. For the positive control, PLA was performed by incubating with primary antibodies (mouse anti-Gag and rabbit anti-biotin) followed by PLA secondary antibodies (anti-mouse IgG coupled to (-) PLA oligo and anti-rabbit IgG coupled to (+) PLA oligo). In the two negative controls, one primary antibody was replaced with a nonimmune control antibody from the same species, as indicated. Addition of other PLA reagents leads to connector oligos linking the (+) and (-) oligos only if the primary antibodies are colocalized; this in turn results in the PLA amplification reaction. Addition of an oligo that recognizes a sequence in the amplified regions and is coupled to a red fluorophore (red star) results in intense red spots at sites where the two primary antibodies are colocalized *in situ*. Thus, red spots indicating colocalization of antigens should be absent when either primary antibody is replaced by a nonimmune antibody. Following PLA, indirect immunofluorescence (IF) is performed by adding a secondary antibody conjugated to a green fluorophore (green star) to detect any unoccupied immune antibody, thus marking Gag-expressing or biotin-containing cells with low-level green fluorescence. (B) A representative field is shown for each of the three antibody pairs and two treatments (the antibodies used are indicated above images, the treatments are indicated to the left). Fields are shown as a merge of three color-channels: the red channel showing biotin-NI PLA, NI-Gag PLA, or biotin-Gag PLA as indicated by red labeling above images; the green channel showing biotin IF or Gag IF as indicated by green labeling above images; and the blue channel showing DAPI-stained nuclei. Scale bars, 10 μm. Graphs below each image column show the average number of PLA spots per cell for each antibody pair and each treatment. Ten to twenty fields were analyzed for each group (containing a total of 107-304 cells per group), with error bars showing SEM.

**FIG. S3.**
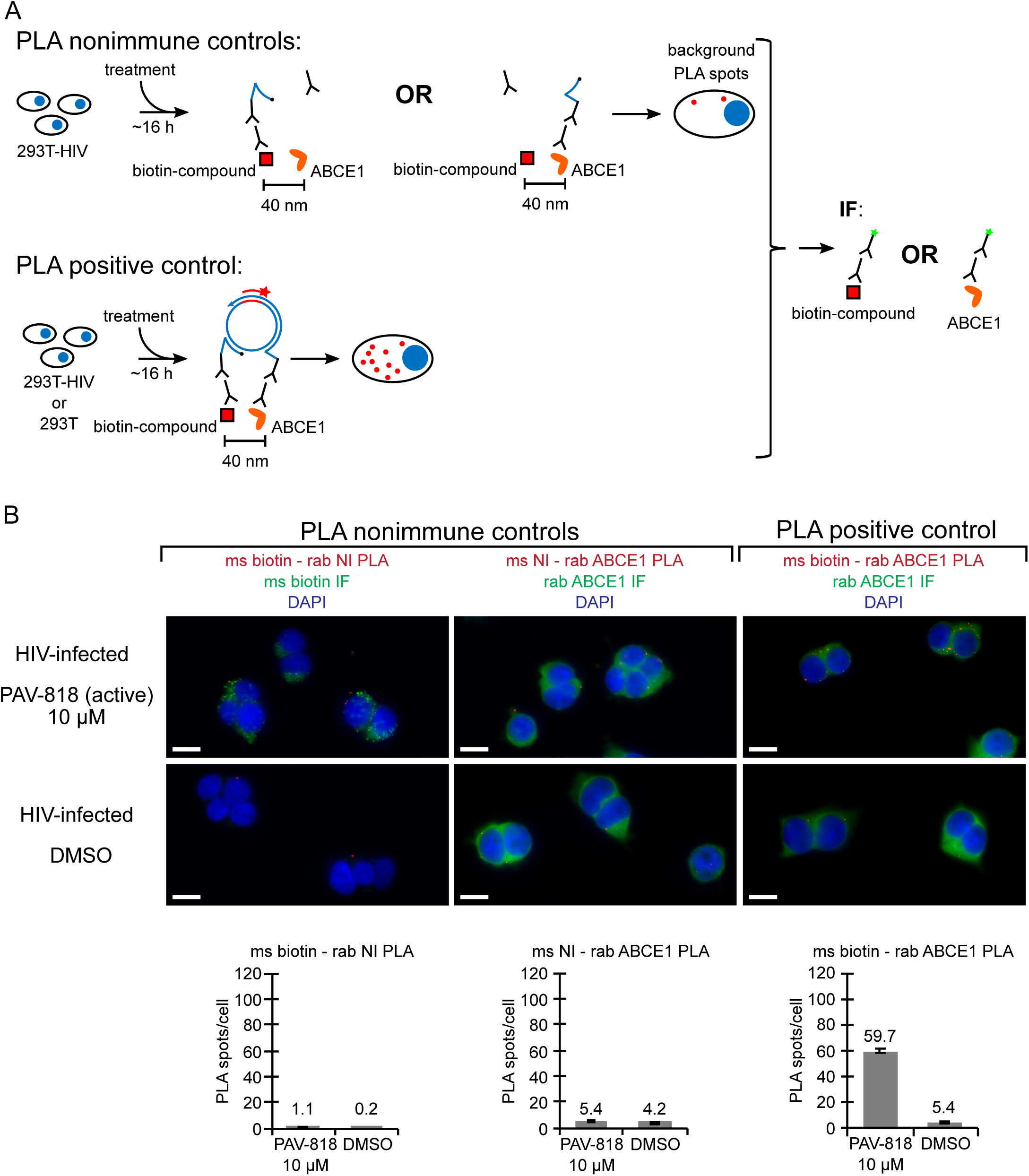
Nonimmune controls for the biotin-ABCE1 proximity ligation assay. (A) Schematic of the PLA approach for testing specificity of the antibodies used to colocalization of compound with ABCE1. 293T cells chronically infected with HIV-1 (293T-HIV) were treated with 10 μM PAV-818 (the biotinylated active compound) or DMSO for 16 h. For the positive control, PLA was performed by incubating with primary antibodies (rabbit anti-ABCE1 and mouse anti-biotin) followed by PLA secondary antibodies (anti-rabbit IgG coupled to (+) PLA oligo and anti-mouse IgG coupled to (-) PLA oligo). In the two negative controls, one primary antibody was replaced with a nonimmune control antibody from the same species, as indicated. Addition of other PLA reagents leads to connector oligos linking the (+) and (-) oligos only if the primary antibodies are colocalized; this in turn results in the PLA amplification reaction. Addition of an oligo that recognizes a sequence in the amplified regions and is coupled to a red fluorophore (red star) results in intense red spots at sites where the two primary antibodies are colocalized *in situ*. Thus, red spots indicating colocalization of antigens should be absent when either primary antibody is replaced by a nonimmune antibody. Following PLA, indirect immunofluorescence (IF) is performed by adding a secondary antibody conjugated to a green fluorophore (green star) to detect any unoccupied immune antibody, thus marking ABCE1- or biotin-expressing cells with low-level green fluorescence. (B) A representative field is shown for each of the three antibody pairs and two treatments (the antibodies used are indicated above images, the treatments are indicated to the left). Fields are shown as a merge of three color-channels: the red channel showing biotin-NI PLA, NI-ABCE1 PLA, or biotin-ABCE1 PLA as indicated by red labeling above images; the green channel showing biotin or ABCE1 IF as indicated by green labeling above images; and the blue channel showing DAPI-stained nuclei. Scale bars, 10 μm. Graphs below each image column show the average number of PLA spots per cell for each antibody pair and each treatment. Five fields were analyzed for each group (containing a total of 44 - 77 cells per group), with error bars showing SEM.

## SUPPLEMENTARY TABLES

**TABLE S1:** **Mutations that arose upon selection in nelfinavir.** Mutations are numbered relative to the HXB2 or NL4-3 genomes, with the frequency of the mutation, the phenotype associated with the mutation, and relevant references indicated.

| Nelfinavir Selection Results |  |  |  |  |  |
| --- | --- | --- | --- | --- | --- |
| Non-synonymous |  |  |  |  |  |
| Appearance in sequencing chromatogram* | Mutation numbered relative to HXB2† | Mutation numbered relative to NL4-3‡ | Frequency‡ of amino acids at mutation position | Mutation associated with: | Notes |
| dominant | **H219Q Gag**<br>(H87Q p24 Gag) | H219Q Gag<br>(H87Q p24 Gag) | H 73.31% (6999)<br>Q 22.86% (2182)<br>P 1.73% (165) | culture adaptation to intracellular CypA level (1,2,3); cyclosporin resistance (4) | in the cyclophilin A binding loop |
| ≥ 50% | **A224T Gag**<br>(A92T p24 Gag) | A224T Gag<br>(A92T p24 Gag) | A 67.49% (6443)<br>P 31.35% (2993) | culture adaptation (5) | in the cyclophilin A binding loop |
| ≥ 50% | **L10F Protease**<br>(L66F Pol) | L10F Protease<br>(L66F Pol) | L 85.68% (4451)<br>I 7.10% (369)<br>V 5.47% (284) | nelfinavir resistance (6); other PI resistance (7) |  |
| dominant | **V77I Protease**<br>(V133I Pol) | V77I Protease<br>(V133I Pol) | V 83.10% (4322)<br>I 16.11% (838) | nelfinavir resistance (8,9) |  |
| dominant | **L90M Protease**<br>(L146M Pol) | L90M Protease<br>(L146M Pol) | L 98.77% (5137)<br>M 1.15% (60) | nelfinavir resistance (6); other PI resistance (7) |  |
| ≥ 50% | **S144R Env**<br>S114R gp120 | S144R Env<br>S114R gp120 | S 24.14% (1248)<br>N 21.88% (1131)<br>T 16.12% (833) | fusion inhibitor resistance (10) | in the Env V1 loop |
| dominant | **A541V Env**<br>A30V gp41 | A539V Env<br>A30V gp41 | A 94.84% (6728)<br>V 2.93% (208)<br>T 1.97% (140) | culture adaptation (11,12); Rev inhibitor resistance (13) | in the Rev response element |
| dominant | **Q550H Env**<br>Q39H gp41 | Q548H Env<br>Q39H gp41 | Q 99.17% (7035) | culture adaptation (11); also arose in PAV-206 selection in current study | in the Rev response element; in the T-20 (Enfuvirtide) drug binding site |

\* Gel chromatograms were visually inspected to identify a second chromatogram peak that is equal or larger in height than the peak corresponding to the reference sequence (defined here as ≥ 50%) or a single chromatogram peak that differs from the reference sequence (defined here as dominant).
† Indicates viral protein to which numbering refers, either in the HXB2 or NL4-3 HIV strain. Numbering of mutation relative to other relevant proteins or domains is indicated in parentheses. Mutations in bold are shown in Fig. 7.
‡ Determined using AnalyzeAlign (available on Los Alamos HIV databases and compendiums site) using the premade Web Alignment; the top three that are > 1% are listed.

**TABLE S2:** **Mutations that arose upon selection in PAV-206.** Mutations are numbered relative to the HXB2 or NL4-3 genomes, with the frequency of the mutation, the phenotype associated with the mutation, and relevant references indicated.

| PAV-206 Selection Results |  |  |  |  |  |
| --- | --- | --- | --- | --- | --- |
| Non-synonymous |  |  |  |  |  |
| Appearance in sequencing chromatogram* | Mutation numbered relative to HXB2† | Mutation numbered relative to NL4-3† | Frequency‡ of amino acids at mutation position | Mutation associated with: | Notes |
| dominant | **V218M Gag**<br>(V86M p24 Gag) | V218M Gag<br>(V86M p24 Gag) | V 87.62% (8365)<br>A 4.85% (463)<br>P 3.40% (325) | culture adaptation to intracellular CypA level (14,3) | in the cyclophilin A binding loop |
| dominant | **N302Y Env**<br>(N272Y gp120) | N300Y Env<br>(N270Y gp120) | N 96.50% (6844)<br>K 1.06% (75) | culture adaptation – CCR5 (15,16,17); fusion inhibitor resistance (18) | in the Env V3 loop; involved in coreceptor binding |
| dominant | **D547G Env**<br>(D36G gp41) | D545G Env<br>(D36G gp41) | G 99.65% (7069) | replication advantage (19,20); fusion inhibitor resistance (21) | in the Rev response element; in the T-20 (Enfuvirtide) drug binding site; D36 in NL4-3 but G36 in most other strains of HIV-1 (19) |
| ≥ 50% | **Q550H Env**<br>(Q39H gp41) | Q548H Env<br>(Q39H gp41) | Q 99.17% (7035) | culture adaptation (11); also arose in nelfinavir selection in the current study | in the Rev response element; in the T-20 (Enfuvirtide) drug binding site |
| Synonymous |  |  |  |  |  |
| ≥ 50% | I<sub>ATT</sub> 205 I<sub>ATC</sub> Pol<br>(I<sub>ATT</sub> 50 I<sub>ATC</sub> Reverse trans.) | I<sub>ATT</sub> 205 I<sub>ATC</sub> Pol<br>(I<sub>ATT</sub> 50 I<sub>ATC</sub> Reverse trans.) |  | unknown |  |
| ≥ 50% | Q<sub>CAG</sub> 552 Q<sub>CAA</sub> Env<br>(Q<sub>CAG</sub> 41 Q<sub>CAA</sub> gp41) | Q<sub>CAG</sub> 550 Q<sub>CAA</sub> Env<br>(Q<sub>CAG</sub> 41 Q<sub>CAA</sub> gp41) |  | replication advantage (22) | in the Rev response element; in the T-20 (Enfuvirtide) drug binding site |
| dominant | V<sub>GTG</sub> 583 V<sub>GTA</sub> Env<br>(V<sub>GTG</sub> 72 V<sub>GTA</sub> gp41) | V<sub>GTG</sub> 581 V<sub>GTA</sub> Env<br>(V<sub>GTG</sub> 72 V<sub>GTA</sub> gp41) |  | fusion inhibitor resistance (23) | in the Rev response element |

\* Gel chromatograms were visually inspected to identify a second chromatogram peak that is equal or larger in height than the peak corresponding to the reference sequence (defined here as ≥ 50%) or a single chromatogram peak that differs from the reference sequence (defined here as dominant).
† Indicates viral protein to which numbering refers, either in the HXB2 or NL4-3 HIV strain. Numbering of mutation relative to other relevant proteins or domains is indicated in parentheses. Mutations in bold are shown in Fig. 7.
‡ Determined using AnalyzeAlign (available on Los Alamos HIV databases and compendiums site) using the premade Web Alignment; the top three that are > 1% are listed.

## SUPPLEMENTARY METHODS

### Synthesis of Tetrahydroisoquinolines

<u>General Procedure for the Synthesis of Substituted Phenethyl amines^a^</u>

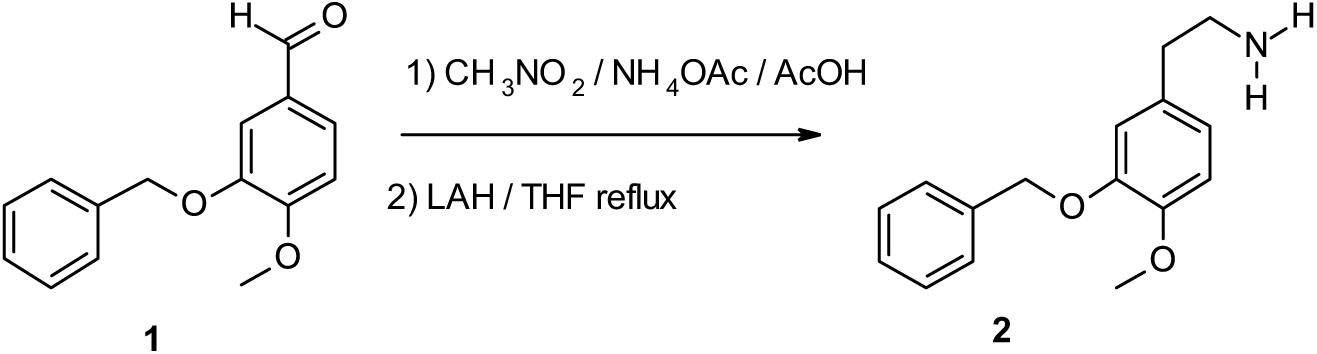

**[0001]** A solution of the Benzaldehyde **1** (0.1 mol), Nitromethane (50 mL, 0.93 mmol) and Ammonium acetate (0.26 mol) in Acetic acid (200 mL) was refluxed for 1 h. Upon cooling the product crystallized out of solution. The crystals were filtered out and washed with a small amount of ether to give a bright yellow nitrostyrene.

**[0002]** To a stirred solution of Lithium aluminum hydride (0.21 mol) in Tetrahydrofuran (270 mL) was added dropwise a solution of the above nitrostyrene (0.11 mol) in Tetrahydrofuran (200 mL). Upon completion the reaction mixture was refluxed with stirring for 16 h, cooled to RT and the excess hydride decomposed by the addition of aq. sat. Na_2_SO_4_. The mixture was filtered and the filtrate rotary evaporated to an amber–brown oil (the title compound) **2** which was used without further purification.

<u>General Procedure for the Synthesis of Substituted Cinammic acids^b^</u>

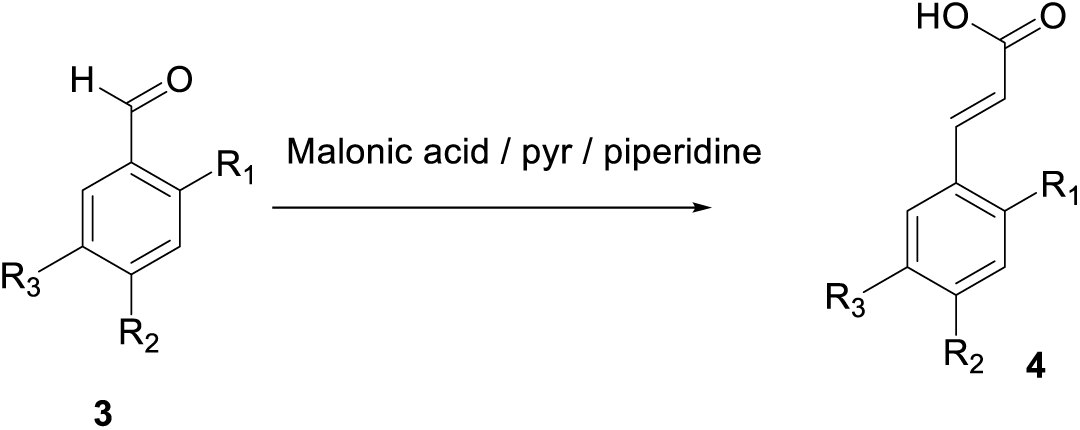

**[0003]** A mixture of the Benzaldehyde **3** (30 mmol), Malonic acid (60 mmol), Pyridine (20 mL) and Piperidine (5 mmol) was stirred at 80°C for 1 h followed by refluxing at 110-115°C for an additional 3 h. The cooled reaction mixture was poured into water (250 mL) and acidified with conc. HCl. The resulting precipitate was filtered off and washed several times with water. It was redissolved in 2M NaOH, diluted with water, acidified with conc. HCl. The solid precipitate was filtered washed several times with water, dried under high vacuum over P_2_O_5_ to afford the titled compound **4**.

<u>General Procedure for the Synthesis of Substituted Cinnamides^c^</u>

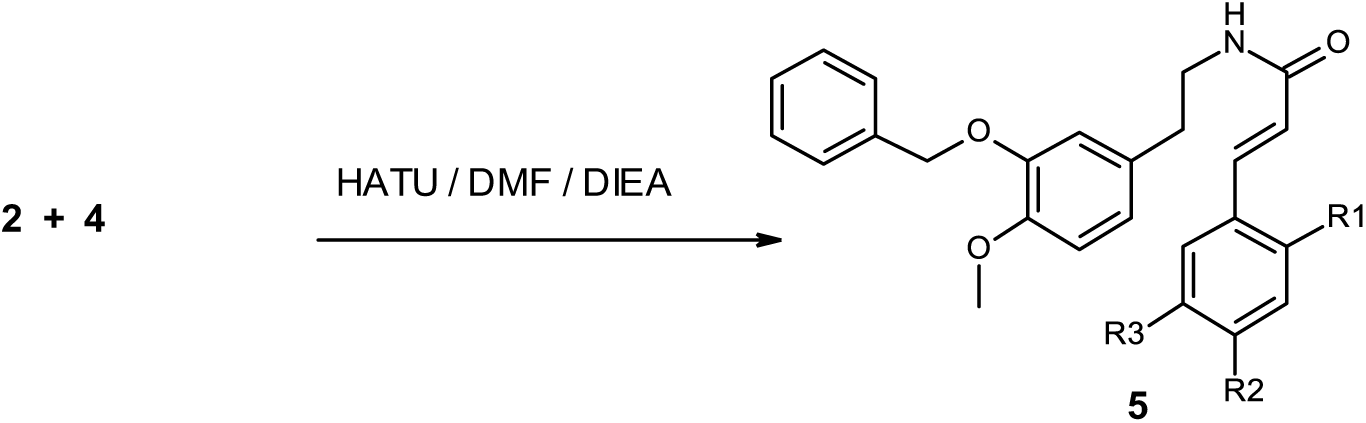

**[0004]** To a stirred mixture of the substituted Phenethylamine **2** (0.48 mmol), substituted Cinnamic acid **4** (0.72 mmol), DIEA [420 μL (2.4 mmol)] & 10 mL of DMF was added HATU (1 mmol). The reaction was stirred at RT for 1 h and then diluted with 20 mL of EtOAc and washed twice with sat NaCl. The EtOAc layer was dried (Na_2_SO_4_) and the solvent removed yielding the title compound **5**.

<u>General Procedure for the Cyclization aCOnd Reduction of the Substituted Cinnamides:</u>

<u>Synthesis of Substituted Tetrahydroisoquinolines^d^</u>

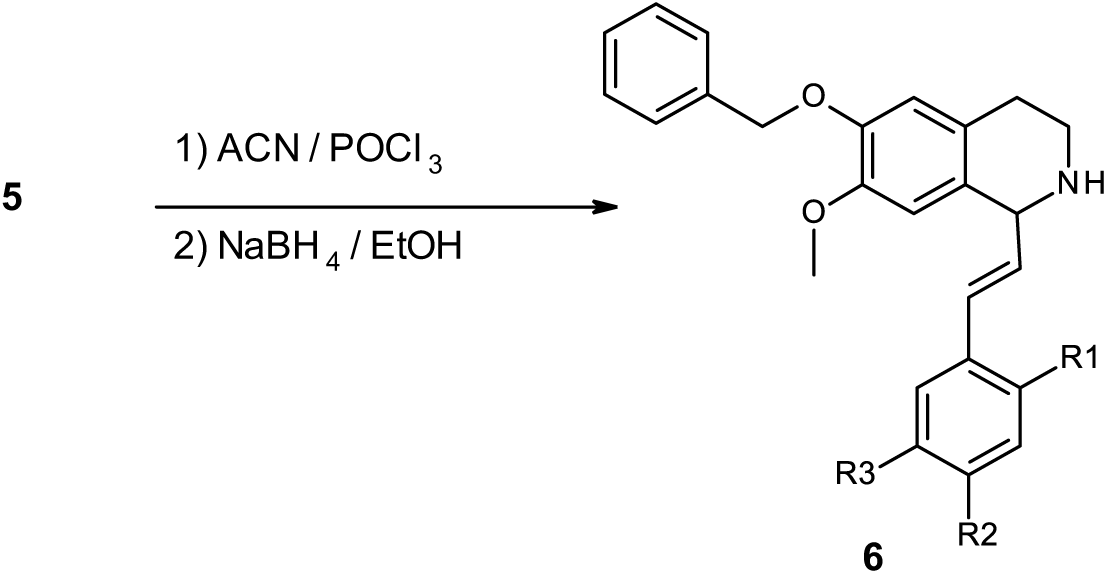

**[0005]** To the substituted Cinnamide **5** (0.563 mmol) in ACN (13 mL) was added, under reflux, POCl_3_ (3.9 mmol). The reaction was stirred at reflux for 30 min and then rotary evaporated to dryness. The residue was taken up into 10 mL of chloroform and was then treated with 20 mL of 2N KOH and 50 mL of Et_2_O. This mixture was rapidly stirred for 30 min at RT and the upper organic layer removed, washed with water, dried (Na_2_SO_4_) and the solvent removed. The resulting dark oil (Substituted Dihydroisoquinoline) was then dissolved into 8 mL of dry EtOH and then treated with NaBH_4_ (0.395 mmol). The excess reagent was destroyed by dropwise addition of 2M HCl, basified with 2M KOH and evaporated to dryness to remove EtOH. The residue obtained was partitioned between water and CHCl_3_, the organic layer was washed with brine, dried (K_2_CO_3_), filtered, concentrated and purified by flash chromatography using 0.4M NH_3_ in MeOH/ CHCl_3_ gradient (and/or purification via RP C18 Prep HPLC using 0.1% TFA Acetonitrile /water) to afford the titled compound **6**.

All compounds were confirmed by Mass Spectrometry:

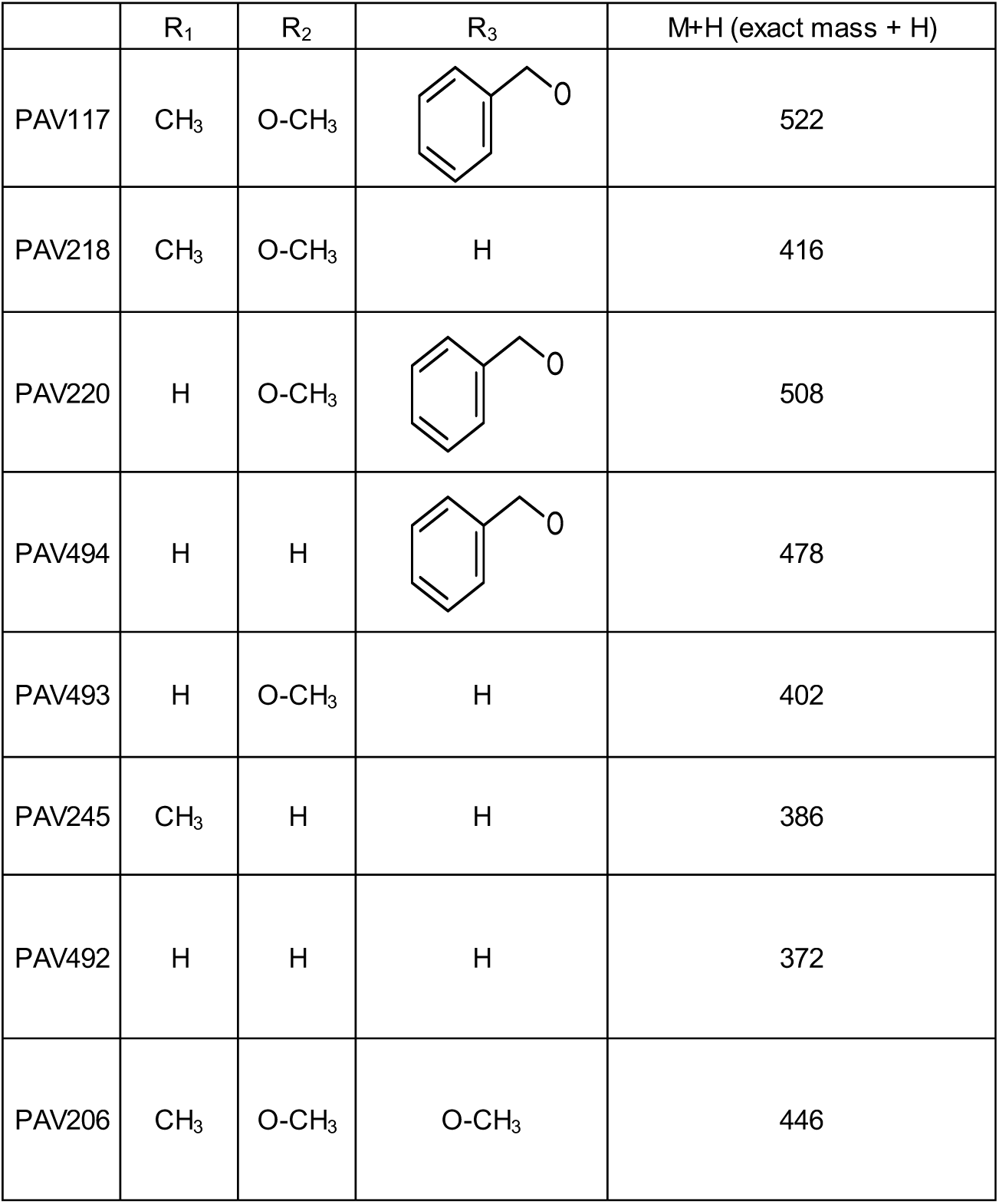

^a^ Cava, M.P.; Buck, K.T. *Tetrahedron* 1969, 25, 2795-2805

^b^ Nag, Ahindra et al *Journal of Molecular Catalysis B: Enzymatic* **2012**, 82, 92-95; Heo, J.N., et al., *Bull. Korean Chem. Soc*., 32 (12), 4431, **2011**; Srikrishna, A. et al, *Synthetic Communications*, 37(6), 965-976; **2007**.

^c^ Patent reference: WO2010001169, Astrazeneca.

^d^ Herbert, Richard B. et al *Tetrahedron* **1990**, 46, 7119-7138.

**<u>Synthesis of N-[2-[2-[3-[[2-[2-[4-[5-[(E)-2-(6-benzyloxy-7-methoxy-1,2,3,4-tetrahydroisoquinolin-1-yl)vinyl]-2-methoxy-4-methyl-phenoxy]butanoylamino]ethylamino]-1-[(3-methyldiazirin-3-yl)methyl]-2-oxo-ethyl]amino]-3-oxo-propoxy]ethoxy]ethyl]-5-[(4S)-2-oxo-1,3,3a,4,6,6a-hexahydrothieno[3,4-d]imidazol-4-yl]pentanamide</u>**

<u>Synthesis of 5-(4-bromobutoxy)-4-methoxy-2-methyl-benzaldehyde:</u>

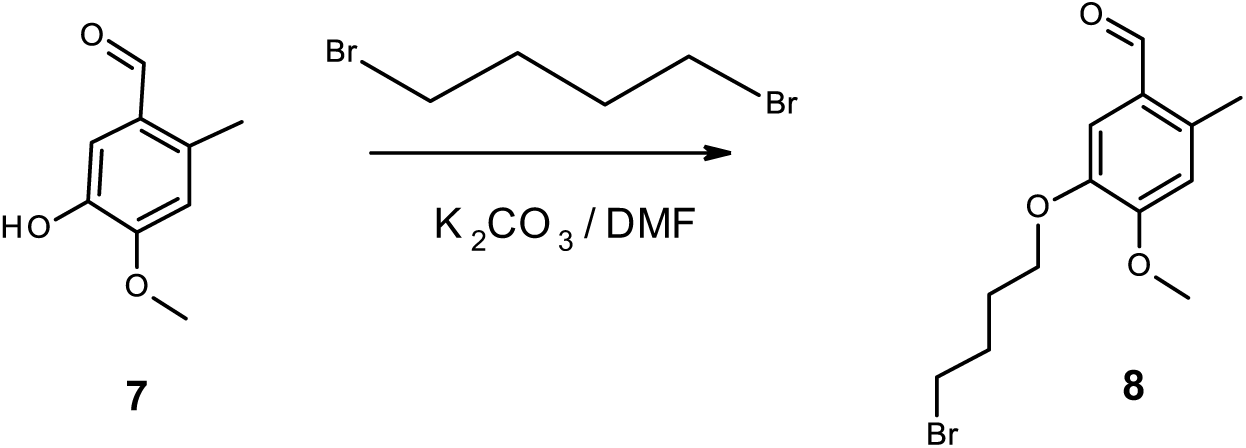

To a solution of Aldehyde **7 [**2.7 g (0.016 mol)] in 30 mL of dry DMF was added K_2_CO_3_ [6.9 g (0.05 mol)]. While stirring at RT under argon atmosphere 1,4-Dibromobutane [10 g (0.048 mol)] was added dropwise. After stirring overnight at RT the reaction mixture was diluted with 100 mL of EtOAc and then washed twice with water. The organic layer was dried (Mg_2_SO_4_), filtered, and rotary evaporated to dryness to afford 4.1 g the desired Bromide **8**.

<u>Synthesis of 5-(3-azidopropoxy)-4-methoxy-2-methyl-benzaldehyde:</u>

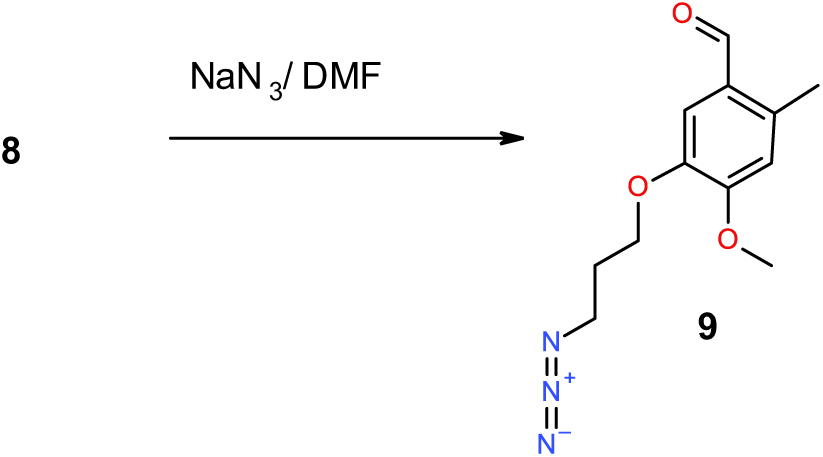

To a solution of Aldehyde **8 [**4.1 g (0.014 mol)] in 30 mL of dry DMF was added NaN_3_ [1.3 g (0.02 mol)]. After stirring at 70°C under argon atmosphere for 5 h the reaction mixture was cooled and then diluted with 100 mL of EtOAc and followed by washing twice with water. The organic layer was dried (Mg_2_SO_4_), filtered, and rotary evaporated to dryness to afford 3.3 g (94%) of the desired Azide **9**.

<u>Synthesis of tert-butyl 1-[(E)-2-[5-(4-azidobutoxy)-4-methoxy-2-methyl-phenyl]vinyl]-6-benzyloxy-7-methoxy-3,4-dihydro-1H-isoquinoline-2-carboxylate:</u>

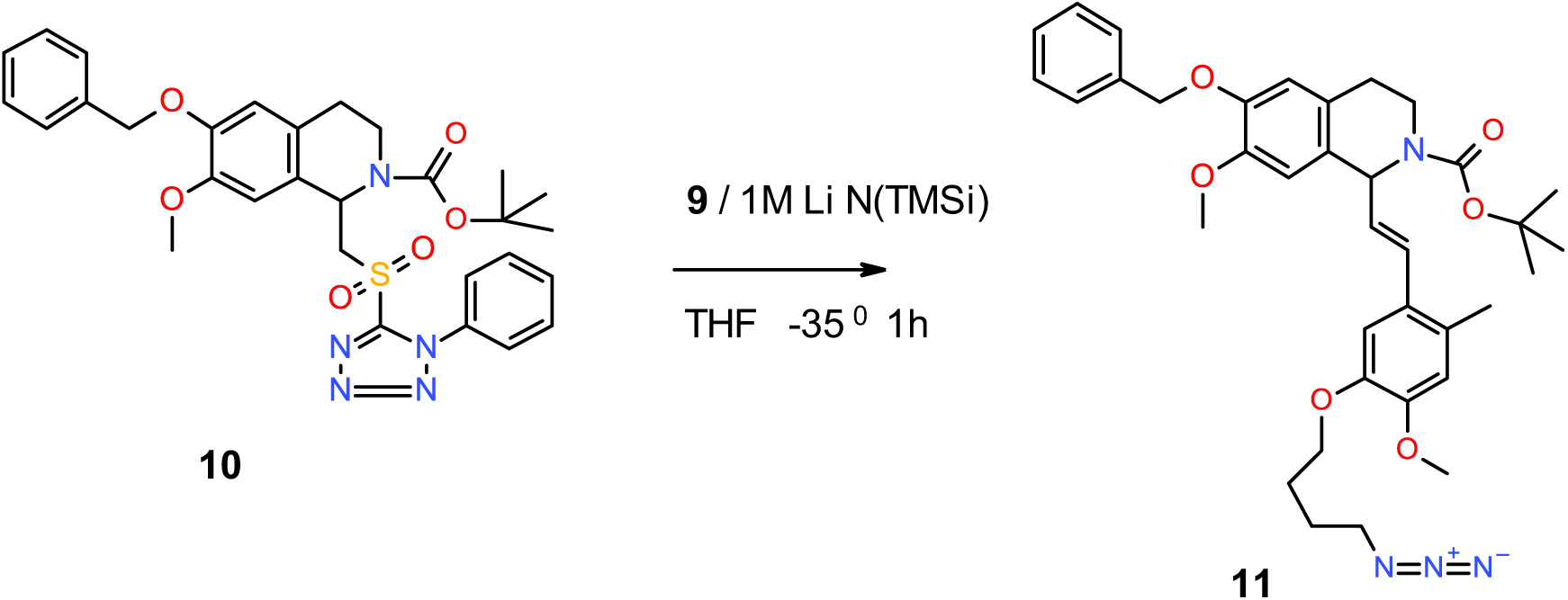

To a solution of Azide **9 [**315 mg (1.2 mmol)] in 8 mL of dry THF was added the Tetrahydroisoquinoline **10**^e^ [236 mg (0.4 mmol)]. The mixture was then cooled and stirred at −35°C under argon atmosphere. Next 1M LiHMDS in THF [1.2 mL (1.2mmol)] was slowly added dropwise and upon complete addition the reaction mixture was stirred at −35°C for 1 h. The reaction mixture was allowed to come to RT and then quenched with 15 mL of saturated NH_4_Cl and then extracted with 20 mL of EtOAc. The organic layer was dried (Mg_2_SO_4_), filtered, and rotary evaporated to dryness to afford the crude Azide **11**. After flash column chromatography using a gradient of EtOAc / Hexane pure **11** (135 mg) was obtained in 54% yield.

<u>Synthesis of tert-butyl 1-[(E)-2-[5-(4-aminobutoxy)-4-methoxy-2-methyl-phenyl]vinyl]-6-benzyloxy-7-methoxy-3,4-dihydro-1H-isoquinoline-2-carboxylate:</u>

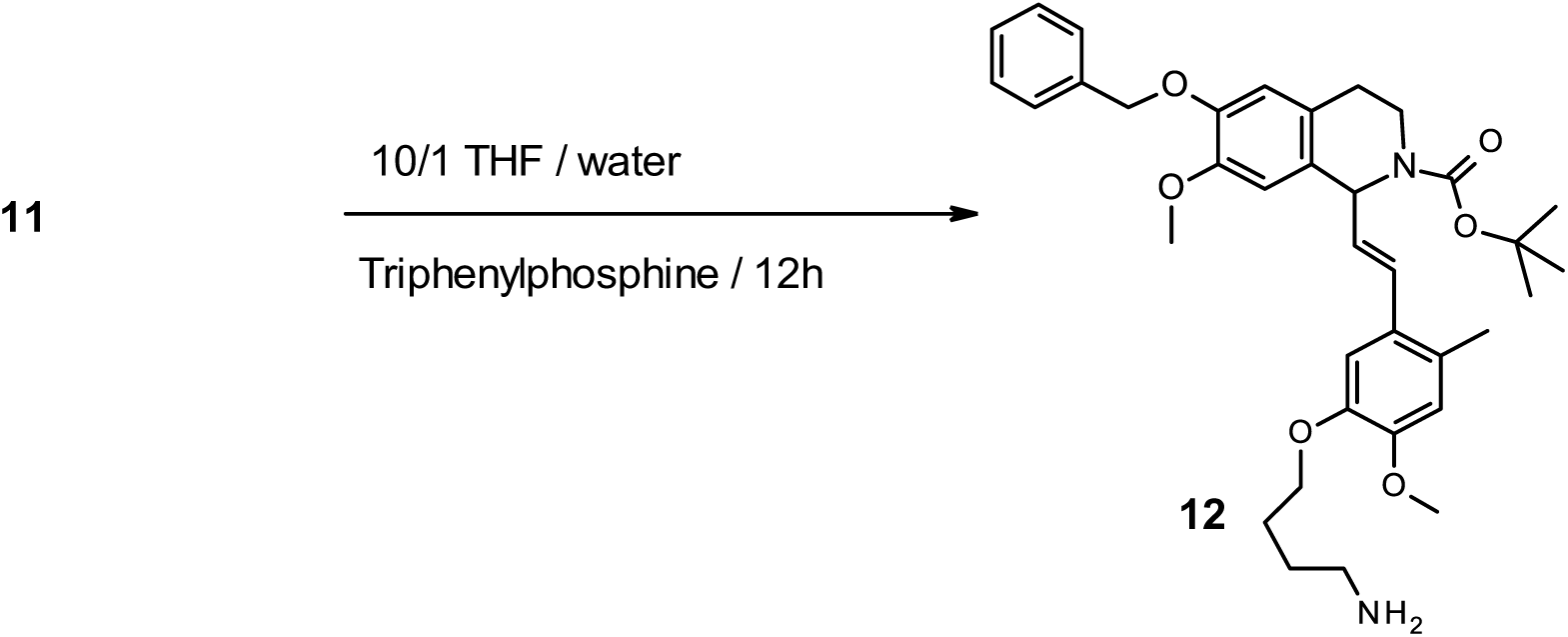

To a solution of Azide **11 [**135 mg (0.215 mmol)] in 5 mL of a 10/1 THF / water was added Triphenylphosphine [64 mg (0.25 mmol)]. The reaction mixture was stirred at RT for 12 h. The reaction mixture was then diluted with 10 mL of water and extracted with 20 mL of EtOAc. The organic layer was dried (Mg_2_SO_4_), filtered, and rotary evaporated to dryness to afford the crude Amine **12**. After flash column chromatography using a gradient of 0.4M NH_3_ in MeOH/ CHCl_3_ pure **12** (90 mg) was obtained in 70% yield.

<u>Synthesis of 3-(3-methyldiazirin-3-yl)-2-[3-[2-[2-[5-[(4S)-2-oxo-1,3,3a,4,6,6a-hexahydrothieno[3,4-d]imidazol-4-yl]pentanoylamino]ethoxy]ethoxy]propanoylamino]propanoic acid:</u>

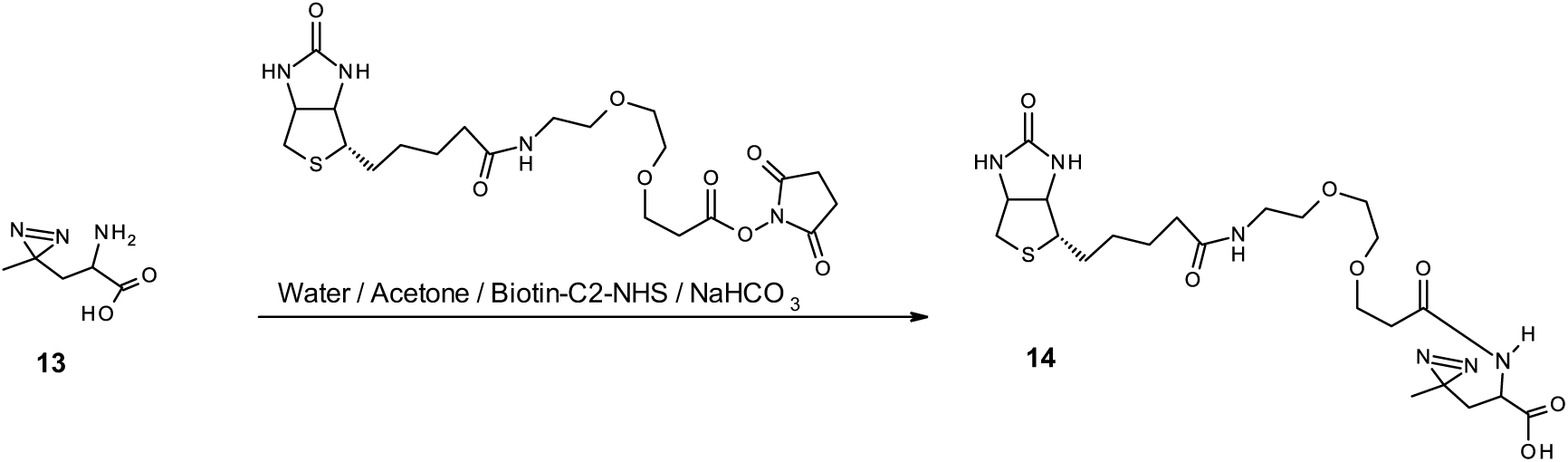

To a mixture of Diazirine **13^f^ [**25 mg (0.17 mmol)], 1 mL Acetone, 1 mL water and NaHCO_3_ [43 mg (0.17 mmol)] was added, with vigorous stirring, Biotin-C2-NHS^f^ [85 mg (0.17 mmol)]. The reaction mixture was stirred at RT for 12 h. The reaction mixture was then diluted with 5 mL of water and extracted with 15 mL of EtOAc. The organic layer was dried (Mg_2_SO_4_), filtered, and rotary evaporated to dryness to afford the crude Acid **14** (40 mg). This was used as was in the next reaction.

<u>Synthesis of N-[2-[2-[3-[[2-[4-[5-[(E)-2-(6-benzyloxy-7-methoxy-1,2,3,4-tetrahydroisoquinolin-1-yl)vinyl]-2-methoxy-4-methyl-phenoxy]butylamino]-1-[(3-methyldiazirin-3-yl)methyl]-2-oxo-ethyl]amino]-3-oxo-propoxy]ethoxy]ethyl]-5-[(4S)-2-oxo-1,3,3a,4,6,6a-hexahydrothieno[3,4-d]imidazol-4-yl]pentanamide:</u>

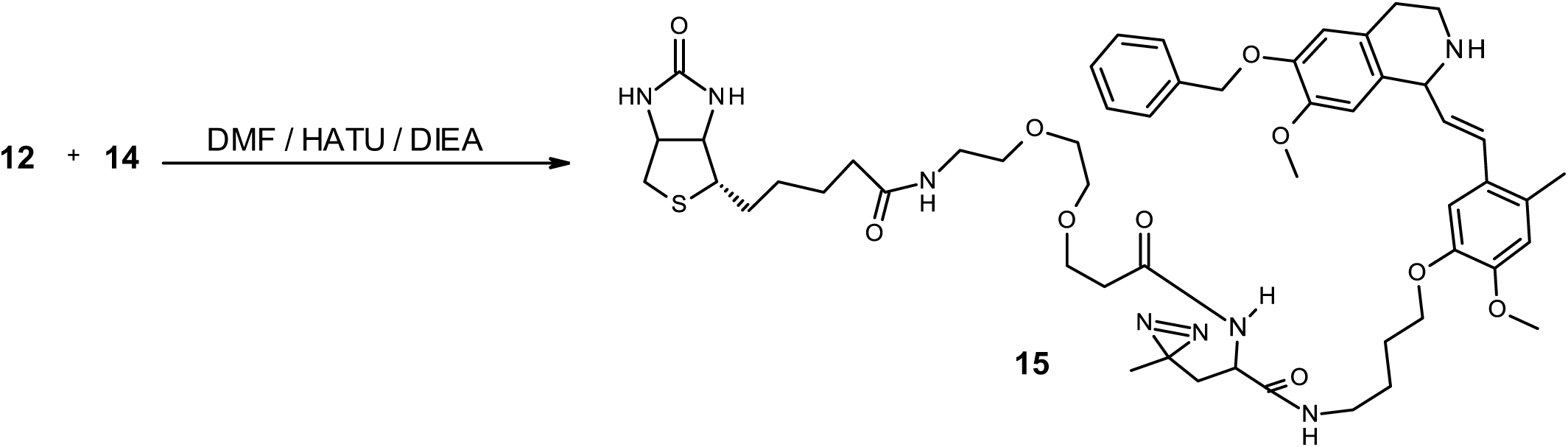

To a mixture of Acid **14 [**40 mg (0.076 mmol)] and Amine **12** [(45 mg (0.075 mmol)] was added 1 mL of dry DMF and DIEA [35 µl (0.2 mmol)]. Next HATU [38 mg (0.1 mmol)] was added in one portion and the resulting mixture was stirred at RT for 0.5 h. The reaction mixture then diluted with 5 mL of EtOAc and followed by washing twice with water. The organic layer was dried (Mg_2_SO_4_), filtered, and rotary evaporated to dryness. The residue was taken up into 1 mL of Acetonitrile / water (90/10) and then purified on a RP C18 Prep HPLC using 0.1% TFA Acetonitrile /water to afford 17 mg (22% overall) of the desired material **15**. M+H=1013

^e^ Reddy, R. et al *Journal of Organic Chemistry* 2012, 77, 11101-11108

^f^ commercially available

